# WormIE: an Integrated Explorer for multi-source *Caenorhabditis elegans* annotations and transcriptomic data interpretation

**DOI:** 10.64898/2026.11.07.764304

**Authors:** Sherlyn P. Wijaya, Ray A. O. Sinurat, Claire E. Richardson

## Abstract

Interpreting gene lists and expression data from *Caenorhabditis elegans* experiments usually means writing scripts, searching scattered databases or uploading unpublished data to a remote server. WormIE (Worm Integrated Explorer, https://wormie.bio) is a free, open-source tool for analyzing userprovided data and exploring *C. elegans* annotations in one browser tab. Starting with gene lists from - omics experiments, genetic screens or other approaches, users can explore what genes have in common and how they interact, test over-representation or ranked-list enrichment in built-in or user-provided gene sets and compare lists by their genes or enriched categories. Of WormIE’s nine collections, which include Gene Ontology and CeNGEN neuron types, two are curated sets found in no other worm resource: neuronal genes and synaptic genes. WormIE counts aligned reads from RNA sequencing and tests for differential expression with the same R package as conventional pipelines, then checks the genes for enrichment in those collections. Clicking any gene in a result page opens the gene page gathering seventeen domains from WormBase, the Alliance of Genome Resources and other sources, such as phenotypes, interactions and expression. Everything runs on the user’s computer without uploading data and works offline. On simulated data, every analysis agreed with its R counterpart (correlation ≥ 0.9999), and in a reanalysis of a germline study of the co-regulator SIN-3, WormIE recovered 99% of the reported differentially expressed genes and all nine WormCat categories. Together, these results show that WormIE reliably links gene lists and expression data to communitycurated knowledge without programming or a server.

## Introduction

For over half a century, the nematode *Caenorhabditis elegans* has been used extensively to connect genes to biology across fields including development, aging, neurobiology and germ cell biology. The knowledge generated from these studies is shared to facilitate ongoing research through a collection of public resources. WormBase (Davis et al. 2022) serves as a central resource for gene models, phenotypes, Gene Ontology (GO) annotations, reagents and literature-based information, while the Alliance of Genome Resources (Agapite et al. 2022) harmonizes orthology, expression and disease data across species. Additional organism-specific resources and tools support more focused uses of these annotations and additional datasets, including CeNGEN (Taylor et al. 2021), which resolves gene expression across the nervous system cell by cell; the WormBase Enrichment Suite (Angeles-Albores et al. 2016; Angeles-Albores et al. 2018), which tests gene lists for enriched tissues, Gene Ontology terms and phenotypes; WormCat (Holdorf et al. 2020), which provides enrichment over a near-complete *C. elegans* genome annotation; and WormEnrichr (Kuleshov et al. 2019), which adapts the Enrichr framework to gene set libraries built from worm-specific resources.

Although these resources are extensive and well curated, some common exploratory workflows could be streamlined. A typical researcher may begin with a single gene, a small candidate list or a set of differentially expressed genes and then ask where those genes are expressed, what phenotypes have been reported, whether they are enriched in a particular cell type or tissue, which biological processes they participate in and which other genes they interact with. Answering these questions can require running analysis scripts that assume computational experience and moving across multiple databases or tools, each with its own interface, identifier conventions and export formats. This creates a practical workflow barrier for users who want to move quickly from a gene or gene list to biological context.

Transcriptomic datasets add an additional workflow barrier because the gene list is not the starting point but the output of an upstream analysis. Established RNA-seq pipelines and tutorials such as Galaxy (The Galaxy Community 2024), iDEP (Ge et al. 2018) and the R-based Bioconductor workflows DESeq2 (Love et al. 2014) and edgeR (Robinson et al. 2010) provide valuable routes through this analysis. Bioconductor workflows run on the user’s own machine and keep data private, which can be an important consideration, but they require programming experience. Web-based tools such as Galaxy and the hosted iDEP server require no programming, but they require uploading the data and depend on the service remaining reachable. Worm-focused expression resources show the value of no-programming interfaces tailored to *Caenorhabditis* transcriptomic data: the *Caenorhabditis* RNA-seq Browser (Akimori et al. 2024) offers expression heatmaps across five *Caenorhabditis* species and differential expression for two of them, and the CaeNDR expression application (Bell and Paaby 2024) opens RNA-seq from 208 wild *C. elegans* strains to gene-by-gene query. Both serve public data curated in advance rather than data the user brings, so users with their own RNA-seq datasets still need separate tools to generate a differential expression list and then additional resources to interpret it. These barriers tend to fall hardest on researchers with limited bioinformatics training, software access, IT support and computational infrastructure, with documented challenges in undergraduate education and resource-limited environments (Williams et al. 2019; Drew et al. 2023; Sangeda et al. 2021), motivating tools that reduce programming, installation and server-dependence requirements.

We present WormIE (Worm Integrated Explorer), a tool that analyzes user-provided gene-level data and carries the results into one integrated database of *C. elegans* annotation. WormIE can test a plain gene list from any source (e.g., RNA-seq, ATAC-seq, proteomics or a genetic screen) for over-representation, shared annotation and interactions among its genes, or compare several lists by their shared genes or enriched terms. Scores and expression matrices open further analyses: ranked-list enrichment, per-sample gene set scoring and two ways to compare ranked-list runs, by leading-edge genes or by consensus. The enrichment tests can also run against the user’s own gene sets in place of the built-in collections. For RNA-seq, WormIE provides the upstream steps: it counts aligned reads, runs differential expression on the counts, compares contrasts with each other and carries the resulting genes into those same analyses, so a gene list can be produced and interpreted in the same place. Clicking a gene in any of these results opens its gene page, which draws on WormBase, the Alliance and other community resources. WormIE links these resources through one stable key, the WormBase gene identifier, so their annotations can be queried together. The resources included were chosen to answer the questions a bench scientist typically asks once a gene is in hand, rather than grouped by where the data came from. The same pages can be browsed directly. WormIE requires no account, no installation, no programming and no paid software. Every search and analysis runs on the user’s own machine, so gene lists and count matrices are not uploaded to a remote server. Once loaded, it also runs offline, which matters where internet connections are slow or costly (Mulder et al. 2016; Boakye Serebour et al. 2024). Here, we describe how WormIE runs, what data it integrates, how its annotations support exploration and which analyses it offers. We also demonstrate that WormIE accurately reproduces a published *C. elegans* analysis, positioning it as a practical companion to WormBase and other established *C. elegans* data resources.

## Methods

### How WormIE works

At its core, WormIE runs in the browser rather than on a server. Opening it downloads the annotation database, one compressed SQLite file of roughly 85 megabytes. Every search, gene page and analysis after that, differential expression included, is computed on the user’s own computer (Figure 1). WormIE can work this way because browsers now run existing C and R programs compiled to WebAssembly (Haas et al. 2017). It uses the same SQLite engine (SQLite Development Team 2025) a local installation would. Through webR (The webR Contributors 2026) it runs DESeq2 (Love et al. 2014) itself, unmodified from Bioconductor (Gentleman et al. 2004; Huber et al. 2015) (Supplementary Note 1). The gene set methods are the exception, since they were reimplemented for the browser and compared with the original packages in *In-browser analysis*. Every analysis is run through a point-and-click interface.

**Figure 1.**
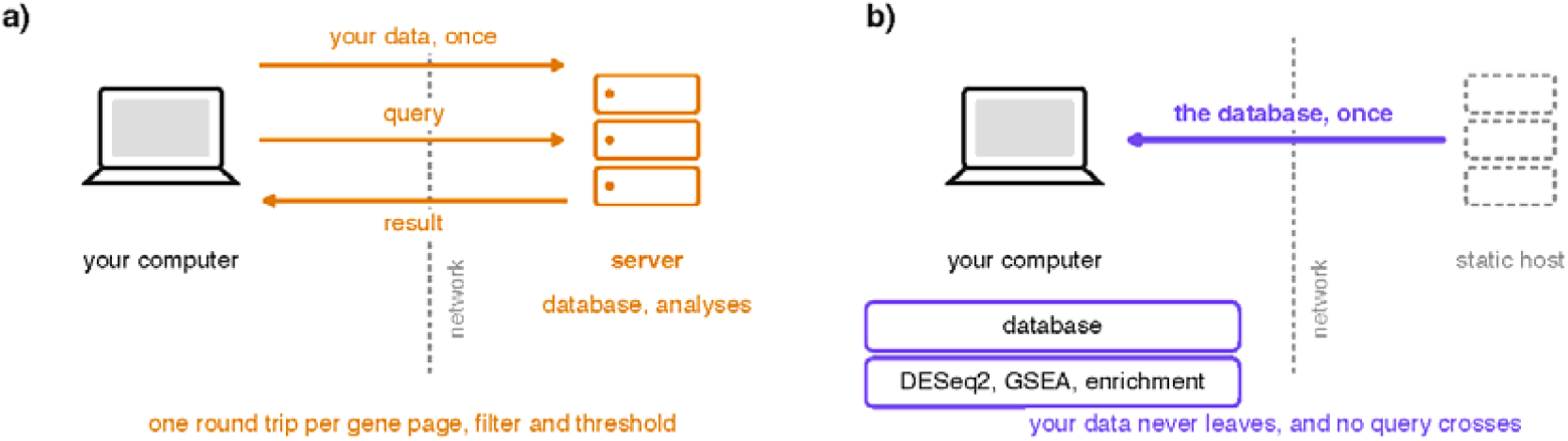
Keeping the database on the user’s computer means no query has to cross the network. Each panel shows one analysis session, with a dashed vertical line marking the network boundary and arrows for everything that crosses it. a) A conventional hosted tool, where the database and the analyses live on the server. The user’s data is uploaded once, after which every gene page, filter and changed threshold is a query sent across the boundary and a result returned, so one session crosses it dozens of times. b) WormIE, where the database crosses once in the opposite direction and nothing is ever uploaded. A newer build is offered when one is published, which happens weekly, and the copy already on the machine stays in use until the user accepts the new one, unless an update to the application needs a newer database structure. The database, differential expression and the gene set analyses all sit on the user’s machine, so every later query is answered without the network.

Downloading everything up front matters because exploration is rarely a single question. A researcher moves from a gene page to a filter to a changed threshold and back again, perhaps dozens of times in one sitting. Where a hosted tool sends each of those steps across the network and waits for the answer (Figure 1a), WormIE answers them from the copy already on the computer (Figure 1b). Every open tab reads that one copy, so a researcher can keep one tab per gene or per comparison without downloading anything again.

Once that first download is done, browsing and gene set analysis no longer need the network. The database is written to the computer’s own storage, so it survives when the tab is closed. Similarly, the programs behind read counting and differential expression are fetched the first time each is used, about 100 megabytes for R and its packages, and then kept the same way. Exporting happens on the computer too: any table is saved as a TSV or CSV file and any plot as an SVG, PNG or PDF file, or all the plots of a page together. WormIE can also be installed from the browser and opened like a desktop application. Installing is optional, but it makes the browser less likely to delete the local copy (Supplementary Note 1). A browser may delete stored files when the computer runs low on space, but it is less likely to do so for a site the user returns to regularly (WHATWG 2026; MDN Web Docs 2026). The database therefore normally stays in place until the user clears the site’s data. Outside well-resourced institutions, where connectivity is often limited or costly, a hosted tool can be unusable rather than merely inconvenient (Mulder et al. 2016; Boakye Serebour et al. 2024). WormIE keeps working in such environments because everything it needs is already on the computer.

The same arrangement keeps a user’s own data on the computer. This protection comes from there being no server to receive the data, not from a privacy policy. For some labs this is the deciding factor rather than a preference, since unpublished data may need to remain on local or institutionally controlled computers. A hosted enrichment tool must receive the gene list or the count matrix before it can answer anything. WormIE requires neither the upload nor a live connection, so it stays usable where policy forbids sending data out or where bandwidth is limited or unreliable.

The worm community has begun to build this way already. ODin (Cho and Sternberg 2026) predicts gene interactions from Alliance orthology entirely in the browser, and its authors attribute the privacy of user data and the tool’s rapid responses to that local processing. Their work also shows that a client-side tool can derive new relationships from curated data rather than only display what is already recorded. Where ODin holds one domain on the machine, WormIE holds all seventeen. The shared-annotation and co-expression panels on each gene page compute new relationships from the local copy in this way (*Exploring the integrated resource*).

### Data ingestion and integration

WormIE compiles seventeen annotation domains. Table 1 lists the public source behind each and the number of records in the build reported here, which uses WormBase release WS298. Each source is authoritative in its own domain, and they rarely name a gene the same way. One gene reaches the build as a WormBase identifier, as a taxon-prefixed protein isoform from STRING (Szklarczyk et al. 2023) and as a numeric Entrez Gene identifier from NCBI (Maglott et al. 2011) and BioGRID (Oughtred et al. 2021). WormIE resolves all of them onto the WormBase identifier and keeps each gene’s public and sequence names alongside its Entrez, UniProt (Bateman et al. 2025), WormPep and GenBank (Sayers et al. 2024) accessions, so a gene can be found by whichever name a user happens to hold. A list pasted from a public dataset and keyed on NCBI gene identifiers therefore resolves to worm annotations without a separate mapping step. Because every record shares that one key, a question can begin at a gene, a phenotype, a tissue or an interaction partner and reach every other domain without leaving the tool.

**Table 1.** Annotation domains integrated by WormIE, the sources they are drawn from, the release of each source and the number of records in the database build. Counts are those of build 20261025-112346, and every build records its own version alongside the release of each source it read.

| Domain | Source | Records | Version / release | Citation |
|---|---|---|---|---|
| Gene models | WormBase | 52,110 genes (19,984 protein coding), 60,156 transcripts | WS298 (final release) | Davis et al. (2022) |
| Genome location | Alliance, WormBase clone map | 46,926 genes placed, 46,890 with a genetic map position | WBcel235 coordinates, WS298 interpolated clone positions | Agapite et al. (2022) |
| Protein domains | WormBase, InterPro | 118,288 assignments (15,693 genes), 9,440 InterPro entries | WS298 | Blum et al. (2025) |
| Phenotype and RNAi | WormBase, Alliance | 119,053 gene-phenotype, 30,181 allele, 90,179 RNAi | WS298, Alliance (unversioned) | Davis et al. (2022) |
| Expression across life stages | WormBase | 324,786 stage medians (46,398 genes, seven stages) | WS298 RNA-seq control libraries | Davis et al. (2022) |
| Tissue and anatomy expression | Alliance, WBbt ontology | 69,328 associations (6,423 genes) | Expression 9.0.0, WBbt of 2026-10-19 (7,191 terms) | Agapite et al. (2022) |
| Protein associations | STRING | 107,389 (combined score ≥ 700) | v12.0 | Szklarczyk et al. (2023) |
| Physical and genetic interactions | BioGRID | 49,089 | 5.0.262 | Oughtred et al. (2021) |
| Functional categories | WormCat | 31,389 assignments, three levels | whole genome v2 | Holdorf et al. (2020) |
| Neuronal genes | Hobert catalog | 1,327 assignments (1,294 genes) | curated for this work | Hobert (2013) |
| Synaptic genes | This work | 178 genes across 12 classes | curated for this work from 139 publications | Supplementary Table S1 |
| Gene Ontology | GO, WormBase GAF | 103,213 direct, 433,011 propagated (12,107 genes) | current.geneontology.org of 2026-07-26 | Ashburner et al. (2000) |
| Orthology | Alliance | 98,628 calls across seven species | Alliance (unversioned) | Agapite et al. (2022) |
| Single-cell neuron types | CeNGEN | 790,889 calls (13,669 genes) at four thresholds | L4 single-cell calls, 2021-02-18 | Taylor et al. (2021) |
| Bulk neuron types | CeNGEN | 327,148 values (52 neuron types) | L4 integrated, contamination-subtracted, 2024-07-17 | Barrett et al. (2026) |
| Disease orthology | Alliance, Disease Ontology | 30,325 associations (4,604 genes) | Alliance (unversioned) | Agapite et al. (2022) |
| Literature | NCBI gene2pubmed, Alliance | 171,125 gene-publication links | not versioned | Sayers et al. (2022) |

One script builds the whole database from those sources ahead of time, retrieving each source without requiring credentials, registration or manual downloads. Most sources are fetched from their distributors as the script runs, while three curated files ship with the build script itself: the WormCat whole-genome annotation (Holdorf et al. 2020) and the neuronal and synaptic gene sets. The neuronal gene catalog follows the categories set out in WormBook’s survey of the neuronal genome (Hobert 2013). The synapse development and function classification was compiled for this work, with an initial literature scan by a language model and every assignment then verified manually, since no maintained resource groups *C. elegans* synapse genes by the role they play. Its twelve classes, the roles within them and the publications behind every gene are listed in Supplementary Table S1. The build reads each curated gene set as a file of gene identifiers and class labels, so a user rebuilding the database can add a curation of their own and use it alongside the rest, both for browsing and for enrichment. After a change to the build script or curated files, a rebuild reuses the inputs already fetched. The weekly rebuild downloads every source again, so the published database does not rely on an outdated local copy.

Beyond fetching its sources, a WormIE build propagates Gene Ontology annotations, calculates a rarity weight for each term, computes gene-gene co-expression across neuron types and creates a full-text search index. Computing these in advance makes WormIE faster: search results appear as a user types, and enrichment tests read prepared tables rather than assembling them during each query.

The propagated annotations change which terms an enrichment test can detect, not just how fast it runs. In the Gene Ontology (Ashburner et al. 2000; The Gene Ontology Consortium 2023), a gene annotated to a specific term also belongs to the broader terms above it. Annotation files, including those of WormBase, record only the most specific term and leave the broader ones to be inferred by the analysis tool that reads them. On the direct annotations alone, genes working in one process through different specific steps would share no term at all, so the process itself might never appear enriched. To prevent this scenario, WormIE computes all of the implied terms during the build, propagating each annotation upward over the same relations that GO.db (Carlson 2025) and clusterProfiler use (Supplementary Note 2). Phenotype and tissue annotations are propagated in the same way, over the is_a links of the phenotype ontology and the is_a and part_of links of the anatomy ontology, as in the WormBase Tissue Enrichment Analysis (Angeles-Albores et al. 2016). Supplementary Note 2 compares WormIE with clusterProfiler (Yu et al. 2012) on a published gene list. Most terms the two tools disagree on trace to the different Gene Ontology releases they read, which is one reason each build records the release date of every source.

Alongside what the build adds, some annotations are deliberately left out. An annotation recording that a gene is not involved in a process stays on the gene page but does not count toward enrichment (Gaudet and Dessimoz 2017). By contrast, phenotype and anatomy annotations of the same kind are not loaded at all, since the interface does not show them. Low-confidence STRING edges and literature links from other species are dropped as well. The exact rules and the reasons for each are given in Supplementary Note 3.

The WormIE annotation database continues to be rebuilt after publication, so each result should name the build that produced the result. Each build records its own version alongside the release or date of every source it read. The About page shows that record for the build in use, with the release, date and license of each source, and the Changelog page lists what changed between builds. Most of those sources continue to change, but WormBase issued WS298 as its final major release in November 2025 and is now maintained as an archive. WormIE’s exported methods text and its BibTeX citation list both name the build used, so a result can be attributed to an exact annotation state. The build can also accept a specified earlier WormBase release, allowing an analysis to use the same WormBase annotations as an earlier run.

### Exploring the integrated resource

WormIE’s analysis tools draw on an integrated collection of annotations organized around WormBase gene identifiers. We first describe how these annotations support the exploration of individual genes and gene lists, then turn to the tools that analyze anything from a gene list to a genome-wide ranking (*In-browser analysis*).

### One search box, every domain

The search box covers the whole database, from gene names, sequence names and identifiers through the curated descriptions to the phenotypes, alleles, tissues, GO terms, WormCat categories, neuronal genes, synaptic genes, protein domains and orthologs, and through a prefix to chromosomes and CeNGEN neuron types (Supplementary Figure S1). A prefix narrows a query to one domain and prefixes combine with AND, OR and NOT, so pheno:locomotion AND expressed:AVA returns the genes with a locomotion phenotype that CeNGEN detects in the AVA neuron type (Supplementary Note 4). A user who has no particular gene in mind can also browse the annotation and reach the genes under each entry (Supplementary Figure S2), starting from eight lists: Gene Ontology terms, WormCat categories, phenotypes, tissues, human diseases, CeNGEN neuron types, neuronal genes and synaptic genes. Protein domains are reached through search instead, since their 9,440 InterPro entries form a flat list too long to scan.

### One gene, one page

Searching for a particular gene leads to its own page, which gathers what is known about it in six groups (Figure 2a). *Identity* opens it, with the gene’s names and accessions, its position on the genetic map and in the genome, its gene model (the isoforms with their exon counts and protein lengths), its protein domains, its orthologs in other species and how widely it is conserved. The map position is given in centimorgans from WormBase’s clone map, and each protein domain links to the list of every gene that carries it. Each accession links out to the matching entry at its source, whether WormBase, the Alliance, UniProt or NCBI. *Perturbation* follows: the alleles available, the phenotypes seen when the gene is knocked down or mutated and the diseases to which it has been linked. *Annotation* lists the gene’s Gene Ontology terms, its WormCat classification from the broad category down to the most specific level WormCat assigns and membership in WormIE’s two curated gene sets (Figure 2b). *Expression* covers where and when the gene has been detected: across tissues, across seven life stages from embryo to adult and dauer, and across neuron types. *Related genes* hold four kinds of partner: experimentally recorded interactions, predicted associations, co-expression across neurons and shared annotation. *Literature* provides two lists: the papers that underlie particular annotations and the wider publication record for the gene. The first is ordered by how many of a gene’s annotations each paper supports, which keeps a claim on the page one click from its evidence (Figure 2c).

**Figure 2.**
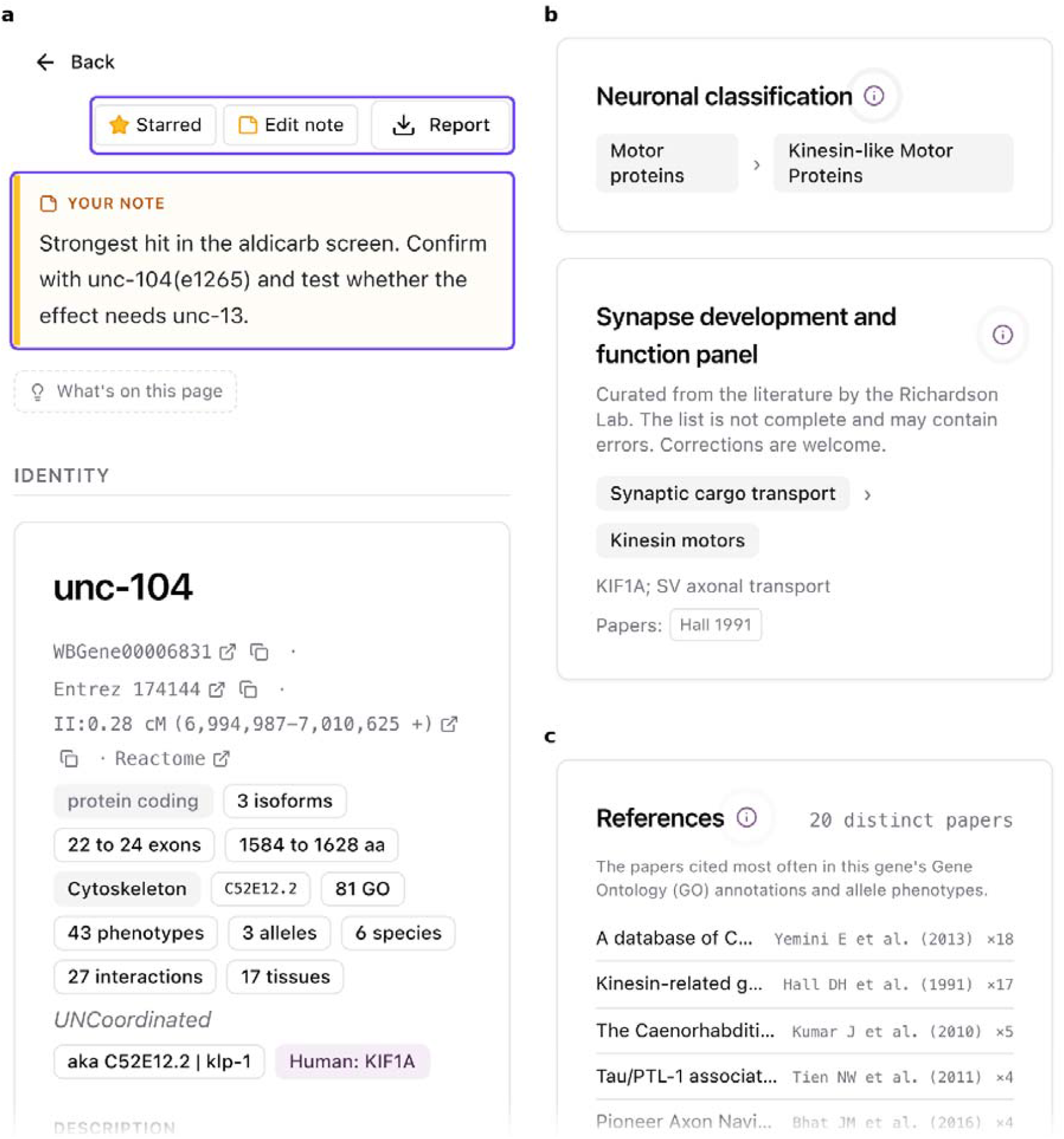
One gene page gathers what is otherwise spread across several portals. a) The top of the page for unc-104. Purple boxes mark the controls for saving the gene, keeping a private note on it and downloading the assembled page as a report. None of these leaves the user’s computer. Below them the page holds six groups of information under their own headings, and only the first group (Identity) is shown here. b) The two curated gene sets, neuronal genes and synaptic genes, which record whether a gene belongs to each, its class and role there and the papers behind the assignment. c) The top of the references list. It holds the papers WormBase cites as evidence for the gene’s Gene Ontology and phenotype annotations, ordered by how many of those annotations each supports. The All literature list further down the page is not shown. It holds every paper NCBI links to the gene with the newest first.

A researcher can write on the gene page as well as read it. They can save a gene, keep a private note of what they have found and download the assembled view as a report (Figure 2a).

### One gene, its neighbors

A standard question asked by a researcher with one or several genes of interest is which genes interact with them. Four panels on the same gene page bring together the evidence relevant to answering this question. The first panel lists the experimentally recorded interactions from BioGRID, and the second lists the predicted associations from STRING. The BioGRID interactions also open as a network that can be widened to the partners of partners and restricted to a neuronal gene category, a WormCat category or a saved list (Supplementary Figure S3). The other two panels rank neighbors by a score WormIE computes, strongest first.

The first of these panels computes similarity from shared GO terms, WormCat categories, neuronal gene categories and interaction partners. GO terms and WormCat categories annotated to fewer genes carry more weight than broad ones. For each suggested neighbor, the panel shows how much the two genes share in each of those areas and names the three most distinctive shared GO terms, so a suggested neighbor arrives with its reason.

The second panel is annotation-independent, comparing the expression profile of the query gene against that of every other gene across the 52 neuron types CeNGEN covers with bulk RNA-seq (Supplementary Figure S4). Two conditions decide whether a gene is kept as a neighbor. First, the correlation between the two profiles must reach a p-value below 0.001, which at 52 observations means a correlation of at least 0.443. Second, each of the two genes must rank among the other’s 200 strongest partners. Broadly expressed genes correlate with much of the genome, so without that second condition they would attach to nearly every list (Supplementary Table S2). Supplementary Note 5 derives both cutoffs and describes the alternative cutoffs rejected in favor of them. The genes surviving both are ordered by mutual rank, the geometric mean of each gene’s rank in the other’s list, so a neighbor ranks highly only when the two genes place each other near the top.

Two genes can correlate in expression without sharing a single annotation, so the co-expression panel can surface candidate interacting genes that an annotation-based search would not. CeNGEN has transcriptionally profiled the nervous system at a depth to which no other tissue has yet been profiled, so every correlation here is taken across neurons. This strategy of correlated expression analysis could be extended to the other tissues when a whole-animal single-cell atlas of comparable depth is made available.

### Many genes, side by side

A list of genes can be checked for how its members interact, compared on what they share and set against other lists. First, though, the names have to agree, because a list may use whichever identifier scheme the analysis pipeline or source literature used. The **Gene name resolver** accepts any mix of established identifiers (gene names, sequence names, aliases, WormBase identifiers or Entrez IDs) and maps each entry to its WormBase identifier. Unmatched entries are gathered separately for review (Supplementary Figure S5). The resolved list can be added to the user’s selected genes, which any page of the tool can use, so the same genes can be browsed, compared and carried into the analyses that follow (*In-browser analysis*) without being pasted again. Gene lists, ranked lists, count tables, fitted models, custom gene sets and whole analysis sessions can also be saved in the browser. The **Datasets** page lists them, reopens a saved session with its results and combines saved lists by union, intersection or difference.

A list can first be checked for how its genes relate to one another. **Network connectivity** counts the interactions recorded between a list’s members, then compares that count with what a random list of the same size would give. The list can be pasted, saved from a search or carried across from an analysis result (*In-browser analysis*). The records come from BioGRID, both physical and genetic by default or either kind alone, rather than from the predicted STRING associations shown alongside them on the gene page (Table 1). STRING draws part of its evidence from co-expression and from text mining, so a list assembled from either would be tested partly against itself. Genes differ widely in how many partners are recorded for them, since a well-studied gene accumulates more. Drawing the random lists from the interaction data at large would therefore make any list of well-known genes look connected, whatever its biology. Each query gene is instead swapped for one carrying a similar number of recorded partners, and a thousand such lists give the distribution the observed count is judged against (Supplementary Note 6). The test then asks whether a list is more connected than the individual prominence of its genes already explains. As an option, the random lists can instead be drawn without considering each gene’s number of recorded partners, so the two results can be compared. The result is a z-score, reported alongside how many of the query genes appear in the interaction data at all. Coverage is uneven, and a list can lose much of its membership before any comparison is made, which is why WormIE prints that second number.

The genes in a list can then be compared for what they have in common. Given a handful of genes, **Annotation overlap** takes the genes side by side, asking which GO terms, phenotypes, tissues, orthologs and neuron-type expression are shared and which belong to only one of them (Supplementary Figure S6). For a pair of genes this is a three-way split: unique to the first, shared, unique to the second. Meanwhile, for three or more it becomes an UpSet plot of the shared and unique terms in each annotation category, a heatmap of pairwise Jaccard similarity and the set of annotations common to every gene in the group.

Whole lists can be set against each other as well. **Gene overlap** takes two or more lists, resolves each the same way and reports which genes are shared and which belong to only one list, an overlap counted on gene identity. A gene that one list holds by its public name and another by its sequence name therefore counts as shared, where a spreadsheet lookup would call it a miss. Any segment of that overlap, the shared hits included, is itself a gene list that can be tested like any other. Whether two lists with few genes in common, such as lists of hits from mutagenesis and RNAi screens, still point to the same biology is a different question, taken up in the *Comparing lists and results*.

### In-browser analysis

The analyses are described in the order a dataset meets them: first the steps from reads to contrasts, then the enrichment analyses, which work on any list or ranking, then the ways to compare lists and results, and last a check of the reimplemented analyses against the original packages (Table 2).

**Table 2.** Analyses available in WormIE, named as the interface names them, with the implementation each uses, the reference method it follows and its agreement with the same analysis run in native R/Bioconductor. Versions are those of the R packages in the WebAssembly analysis bundle, built against R 4.5.1, that ships with the database build reported in Table 1. Analyses computed directly in the browser, rather than through the R bundle, carry no package version.

| **Analysis** | **Implementation** | **Version** | **Follows** | **Agreement** |
|---|---|---|---|---|
| Read counting | featureCounts compiled to WebAssembly | subread 2.0.6 | Liao et al. (2014) | same program |
| Differential expression | DESeq2 in webR, with apeglm shrinkage, or ashr where the baseline is not the reference | DESeq2 1.52.0, apeglm 1.34.0, ashr 2.2-63 | Love et al. (2014) | r = 1.0000, no sign flips |
| Contrast comparison | two contrast result tables compared directly | n/a | no reference implementation | not applicable |
| Over-representation | reimplemented, not loaded from R | n/a | Yu et al. (2012) | 121 of 121 sets |
| Ranked-list enrichment | reimplemented, not loaded from R | n/a | Subramanian et al. (2005) | r ≥ 0.9999, 114 of 114 sets |
| Custom gene set testing | either of those two tests run against a user-supplied GMT file | n/a | as over-representation and ranked-list enrichment | covered by those rows |
| Network connectivity | interaction count against a degree-matched permutation null | n/a | Rossin et al. (2011) | no reference implementation |
| Enrichment overlap | over-representation run on each list, the term tables then compared | n/a | no reference implementation | covered by the over-representation row |
| Leading-edge overlap | leading-edge genes of saved ranked-list runs compared as gene lists | n/a | no reference implementation | covered by the ranked-list row |
| Ranked-list consensus | sets significant in several saved ranked-list runs with overlapping leading edges | n/a | no reference implementation | covered by the ranked-list row |
| Per-sample gene set scoring | reimplemented, not loaded from R | n/a | Barbie et al. (2009) | r = 1.0000 |
| Trend test on gene set scores | limma in webR, one linear model per set | limma 3.68.5 | Hänzelmann et al. (2013) | same package |

### From reads to contrasts

Analysis of RNA-seq data can begin from aligned reads in BAM format or from a count matrix the user already has. Only a BAM file needs **Read counting**, which summarizes its reads into a per-gene count matrix in the browser using featureCounts (Liao et al. 2014) compiled to WebAssembly. With one click, it counts against the gene annotation of the WormBase release the database was built from. This step is simple because WormIE serves only *C. elegans*, so there is no organism, genome or annotation to choose before counting. It also takes an annotation file of the user’s own in GTF or SAF format, so reads can be summarized on the same release as a published analysis rather than on the current one. Reads are counted as unstranded, stranded or reversely stranded to match the library preparation. Differential expression follows in the same session, and its results feed two further analyses: over-representation as a filtered list of differentially expressed genes and ranked-list enrichment as every gene ranked by its test statistic, which combines the size and the certainty of its change. A differential expression table computed elsewhere, for example in edgeR (Robinson et al. 2010), limma (Ritchie et al. 2015) or PyDESeq2 (Muzellec et al. 2023), can be pasted in at that same point, so a laboratory with a preferred pipeline reaches the worm annotation without rerunning the comparison here. Counts, differential expression and then enrichment is the usual route through RNA-seq data (Conesa et al. 2016), and WormIE follows it from reads to enrichment. Along that route, clicking a gene in a plot of the results or in an enriched set opens its gene page (*Exploring the integrated resource*), so a result leads directly to what is known about that gene.

**Differential expression** runs DESeq2 (Love et al. 2014) itself, the same package a local R analysis would use, rather than a reimplementation of it. DESeq2 fits its model to the whole count matrix, estimating for each gene how much it varies between replicates. WormIE keeps that fit alongside two other versions of the counts: normalized and variance-stabilized (Anders and Huber 2010). The normalized version corrects for sequencing depth so a gene reads comparably across samples. A single gene’s expression is therefore plotted from the normalized counts. The sample PCA plot and the sample-distance heatmap use the variance-stabilized counts instead, which put every gene on roughly the same scale of variation (Supplementary Note 7). Variance stabilization goes no further than those views, and every analysis that reads an expression matrix reads the normalized version.

A contrast is then extracted from the DESeq2 fit rather than from a new one, so the model is fitted once regardless of the number of comparisons. Each contrast gives every gene a log fold change that represents how far its expression moved between the two conditions. Fold changes are shrunk with apeglm (Zhu et al. 2019), the DESeq2 default. Because apeglm shrinks only comparisons made against the reference condition, a comparison whose baseline is another condition is shrunk with ashr (Stephens 2017) instead (Supplementary Note 7). Both are defaults rather than fixed choices, since the user can pick apeglm, ashr or no shrinkage. The case study turns shrinkage off to match a published analysis (*A published analysis, reproduced in a browser*).

WormIE asks which condition each sample belongs to, such as genotype or treatment, optionally alongside a batch or replicate effect. Any number of conditions can enter the model, and each comparison is pairwise. For instance, a WormIE user can fit wild type and two mutants in one model, then compare each mutant against the wild type in its own table. Each comparison is shown as a volcano plot, an MA plot and a p-value histogram above a filterable gene table. For a trajectory, or for any question spanning more than two conditions at once, the fit can be fed to a different analysis: the per-sample gene set scoring described in the *Enrichment analyses*. For any analyses not available in WormIE, the fitted DESeq2 object can be downloaded and the analysis continued in R without recomputing the fit, or uploaded again later to extract further contrasts without refitting. A result table produced outside WormIE can then be pasted back in, so a design that WormIE does not fit itself can still be assessed with all the analyses downstream of the contrast.

Two contrasts can also be set against each other in **Contrast comparison**, which compares the full result tables rather than only the significant genes drawn from them. A typical pair is the same change measured in two genetic backgrounds, such as aging in wild type and aging in a mutant, and the question is whether the two contrasts move the same genes in the same direction and which genes they disagree on. Every gene present in both tables falls into exactly one group at an adjusted p-value below a cutoff the user sets, 0.05 by default. Three groups hold the genes significant in both contrasts (up in both, down in both or flipped, meaning up in one and down in the other), and the rest are significant in one contrast alone or in neither. A Venn diagram of the two significant sets counts the three shared groups in its overlap and divides each contrast’s own region by direction. WormIE also draws a scatter plot of the log fold changes of one contrast against the other, one point per gene, colored by the three shared groups, and reports the Pearson correlation of the log fold changes over the genes present in both. Each quadrant of the plot is labeled with the share of its genes that fall in the matching group, so the fraction of genes that rose in both contrasts, for instance, can be read off directly. Each group can be inspected on its own, and since a point in the plot opens that gene’s page, any gene the two contrasts disagree on can be looked up directly. The comparison is descriptive, and whether a mutation changes how genes respond is formally a test of the interaction between the two factors, which the downloaded DESeq2 fit allows in R.

### Enrichment analyses

A contrast from the previous step leads straight into the enrichment analyses, which also accept a plain gene list, a list of genes with scores or an expression matrix from any source.

**Over-representation** takes the genes of a list (worm genes, or human, mouse, rat or zebrafish genes via their worm orthologs), such as those a contrast marked as significantly changed, and asks whether an annotation turns up among them more often than chance would give. Here, significant means an adjusted p-value below 0.05 by default, and any fold-change requirement is set by the user. If differential expression ran in WormIE, the contrast table carries straight over, already filtered on that standard. A results table produced elsewhere, by clusterProfiler, GSEApy or topGO, can be loaded as well and drawn with the same plots.

The annotations tested come from nine collections: Gene Ontology, WormCat category, phenotype, tissue, human disease, protein domain (InterPro), neuron type (CeNGEN), curated neuronal genes and curated synaptic genes. The two curated collections, neuronal genes and synaptic genes, are found in no other worm resource, and a result from each is shown in Supplementary Figure S7. Every collection is tested the same way, with a hypergeometric test, so choosing one changes what the query genes are compared against rather than how the comparison is made. The universe for each collection is the set of genes annotated in it, and a query list is reduced to its annotated members before testing (for Gene Ontology, one aspect at a time, Supplementary Note 2). A gene that is not associated with any term in that collection is therefore left out of both the query and the universe.

Apart from the propagation of the Gene Ontology, phenotype and tissue annotations (*Data ingestion and integration*), the gene sets come directly from the source, with two exceptions. Neuron-type enrichment is the first. CeNGEN data are expression levels of each gene in each neuron type, whereas a hypergeometric test needs a yes-or-no answer for whether a gene belongs to a set. WormIE therefore uses CeNGEN’s own thresholding (Taylor et al. 2021) to decide which genes are expressed in each neuron type and builds a gene list for every type (Supplementary Note 3), and Supplementary Figure S8 shows one result. At the threshold used, CeNGEN estimates a true positive detection rate of 0.81 and a false discovery rate of 0.14, with the detection bar set gene by gene rather than once for all. WormCat is the second. It assigns each gene a category at three nested levels, and WormIE tests all three in one ranked list rather than in the three separate outputs WormCat produces (Holdorf et al. 2020). The ranked list therefore shows the level of description at which a signal sits. A category that WormCat repeats unchanged at the next level is tested once. The cost is that a broad category and the narrower ones beneath it, overlapping by construction, share one multiple-testing correction. WormIE accepts that trade to keep WormCat consistent with the other collections, each of which produces one ranked list under a single correction (one for each aspect of Gene Ontology).

A pasted list is tested against every gene annotated in the collection by default. A list taken from a contrast is tested instead against the genes in that contrast’s results table, since a gene filtered out before testing could never have been called differentially expressed. The user can replace either background. For instance, in tissue-specific RNA-seq a gene not expressed in the profiled tissue could never have appeared among the hits. Once the user supplies every gene expressed in that tissue as the background, WormIE recomputes the universe and the per-term counts from those genes. Only terms holding at least two query genes are tested. They are ranked by p-value and reported with a Benjamini-Hochberg adjusted value (Benjamini and Hochberg 1995) computed across those terms. Because GO terms are nested, many enriched terms in one list are near relatives of each other, and a specific term and the broader terms above it often pass the test on the same genes. A plot of the top twenty-five terms can then spend most of its rows on one signal. To get more out of the same output, WormIE implements a reduction that follows the REVIGO procedure (Supek et al. 2011). In short, REVIGO takes the two most similar terms in the list, keeps one by its own rules (a very broad term is folded first, and otherwise the term with the smaller p-value is kept) and folds the other under it, then repeats with the next most similar pair until no two remaining terms are more similar than a cutoff the user sets. Hence a higher cutoff tolerates more similarity and folds fewer terms, and a lower cutoff gives a shorter list of more distinct terms. The default of 0.7 is REVIGO’s own. Folded terms are not removed but only hidden under their representative, so the full list is one click away and no p-value changes. WormIE applies the same rule to the other collections by measuring similarity as the overlap of the query genes behind two terms instead, with a default cutoff of 0.5 (Supplementary Note 8). The reduction is switched off by default and applies to the table and the figures alike.

As an alternative to over-representation, **Ranked-list enrichment** keeps every expressed gene rather than only those that changed. It follows the preranked gene set enrichment analysis (GSEA) method of Subramanian (Subramanian et al. 2005), reimplemented for the browser after the faster algorithms of fgsea (Korotkevich et al. 2021). It works on any list of genes, as long as each gene carries a score, since names alone carry no information on how each gene changed in the sample. A list carried over from WormIE’s differential expression step is ranked by its Wald statistic. A pasted results table is ranked by its signed test statistic where it has one, and by −log10 of its p-value signed by the fold change where it does not. A list of genes with scores of the user’s own, such as fold changes, is ranked by those scores.

The gene sets come from the same nine collections that over-representation uses. For example, the genes assigned to one GO term or one curated class are taken as a set. Similarly, a user’s own gene sets can be used for over-representation or ranked-list enrichment through **Custom gene set testing**. Sets with fewer than 15 or more than 500 genes are left out by default, the defaults of the original GSEA software. Each set is scored in turn by walking down the full ranked list and keeping a running total. The total rises at each gene belonging to the set by an amount reflecting how strongly that gene moved, and it falls by a fixed amount at each gene that does not. The score is how far that running total strays from zero, so it is highest when the set’s genes crowd the most upregulated end of the list and most negative when they crowd the most downregulated. The set’s members between that end of the list and the point where the total peaks are its leading edge, the genes that produced the score.

That score is then compared with the scores of a thousand random gene sets of about the same size, drawn from the full ranked list (sets within roughly 10% of each other in size share one collection of random sets). The p-value is the proportion of those random sets with scores at least as extreme in the same direction. A set that beats all or nearly all of those thousand can be given no p-value below 1 in 1,000, the limit of the method, so the strongest sets finish tied or nearly so. To separate them, WormIE re-estimates the p-value of every set within three times that limit by fgsea’s multilevel method (Korotkevich et al. 2021), which reaches far smaller values, down to the floor of 10^−50^ that fgsea also uses. The p-value counted from the thousand random sets remains available beside the refined estimate, so the two can always be compared.

A time course sampling one strain across development has expression data, but its question is the trajectory rather than any one pair of timepoints. **Per-sample gene set scoring** answers that question, and it is the one enrichment analysis that takes an expression matrix rather than a gene list. It scores each sample separately against every gene set in a chosen collection, either one of the collections the other analyses use or a GMT file of the user’s own, following the ssGSEA method (Barbie et al. 2009) as GSVA implements it (Hänzelmann et al. 2013). A built-in set is scored by default only when 15 to 500 of its genes are present in the matrix (Supplementary Note 9). Users can paste in an expression matrix or carry one over from differential expression, which hands over the normalized counts for every sample in the fit. In a time course, this produces a score for each gene set in every replicate at every timepoint, so a trend against a continuous variable such as age or dose is visible in the scores themselves. The scores are descriptive rather than a test, so a claim about a trend or a difference between conditions is made by a second step that fits a linear model to each set’s scores across the samples with limma in the browser (Supplementary Note 9). Scoring each sample also keeps replicates visible, so a sample that behaves unlike the rest of its group can be identified.

### Comparing lists and results

Gene overlap and Annotation overlap, described in the *Exploring the integrated resource*, compare lists on their genes and annotations. The three analyses here compare what the lists enrich, a question that matters when two lists share few genes.

**Enrichment overlap** takes more than one gene list and compares them on the terms enriched in each rather than on their members. This can reveal shared biological functions even when the lists contain few of the same genes. Over-representation runs on each list separately, and the resulting term tables are compared the way Annotation overlap compares genes in the *Exploring the integrated resource*, with an UpSet plot and a heatmap of pairwise Jaccard similarity showing which functions reach significance in which list.

**Leading-edge overlap** takes two or more saved ranked-list results and compares them on their leading-edge genes rather than on the names of their enriched sets. Each result contributes the pooled leading edge of its significant sets, and the pooled lists are compared with the same UpSet plot. This shows whether two analyses are driven by the same genes even when the sets they enrich differ in name.

**Ranked-list consensus** takes the same saved results and lists the sets that reach significance in at least a chosen number of them. A set must also keep much the same leading edge across those results, measured by a Jaccard index above a threshold the user sets, so a set enriched several times for different genes each time is left out. The survivors are ranked by how strongly and how consistently they are enriched, so a set that recurs with the same genes outranks a single large hit.

## Results

### The results match the original packages

Over-representation, ranked-list enrichment and per-sample gene set scoring are the three analyses WormIE reimplements, so we validated them by comparing their results with those from the reference packages in native R/Bioconductor (R Core Team 2025; Gentleman et al. 2004) (Table 2, Supplementary Note 10).

WormIE and R ran on the same synthetic inputs, generated from a fixed seed, with a signal planted for the significance tests so that some sets reach significance. Results from WormIE closely matched those from the reference R packages. Over-representation identified all 121 of R’s significant sets and added none. Ranked-list enrichment scores correlated at r ≥ 0.9999. For the strongest sets, whose p-values come within three times the 1-in-1,000 limit of the random sets, both tools refine the estimate, and the refined p-values agreed at r > 0.998. With those refined estimates, WormIE recovered all 114 sets that fgsea called significant and added none. Per-sample gene set scores correlated at r = 1.0000, with no score differing by more than 0.0001. The few differences in raw p-values between WormIE and R occurred among sets near the significance threshold. Supplementary Note 10 accounts for each, and gives the timings, the hardware and the full per-analysis tables.

WormIE runs DESeq2 directly rather than reimplementing it, but we also tested whether its WebAssembly version produced the same results as R. The WormIE and native R runs correlated at r = 1.0000 on log fold changes, and no differentially expressed gene changes the sign of its log fold change.

Table 2 gives the package versions these numbers were produced with. A later build may carry newer ones. In the case study, two DESeq2 releases eight Bioconductor cycles apart gave the same calls on the same input (*A published analysis, reproduced in a browser*).

### A published analysis, reproduced in a browser

To test whether an analysis run in WormIE reaches the same conclusions as one run conventionally, we reanalyzed data from Robert et al. (2023). That study asked how the transcriptional co-regulator SIN-3 shapes the germline transcriptional program, comparing dissected germlines from *C. elegans* wild type against the *sin-3(syb2172)* deletion by RNA-seq. We retrieved their six libraries of raw reads, three per genotype, from the Sequence Read Archive (BioProject PRJNA944713, GEO series GSE227368), and aligned them outside WormIE with the aligner the published analysis used. The rest of the analysis was then repeated in WormIE, including read counting, differential gene expression and enrichment analysis (Figure 3).

**Figure 3.**
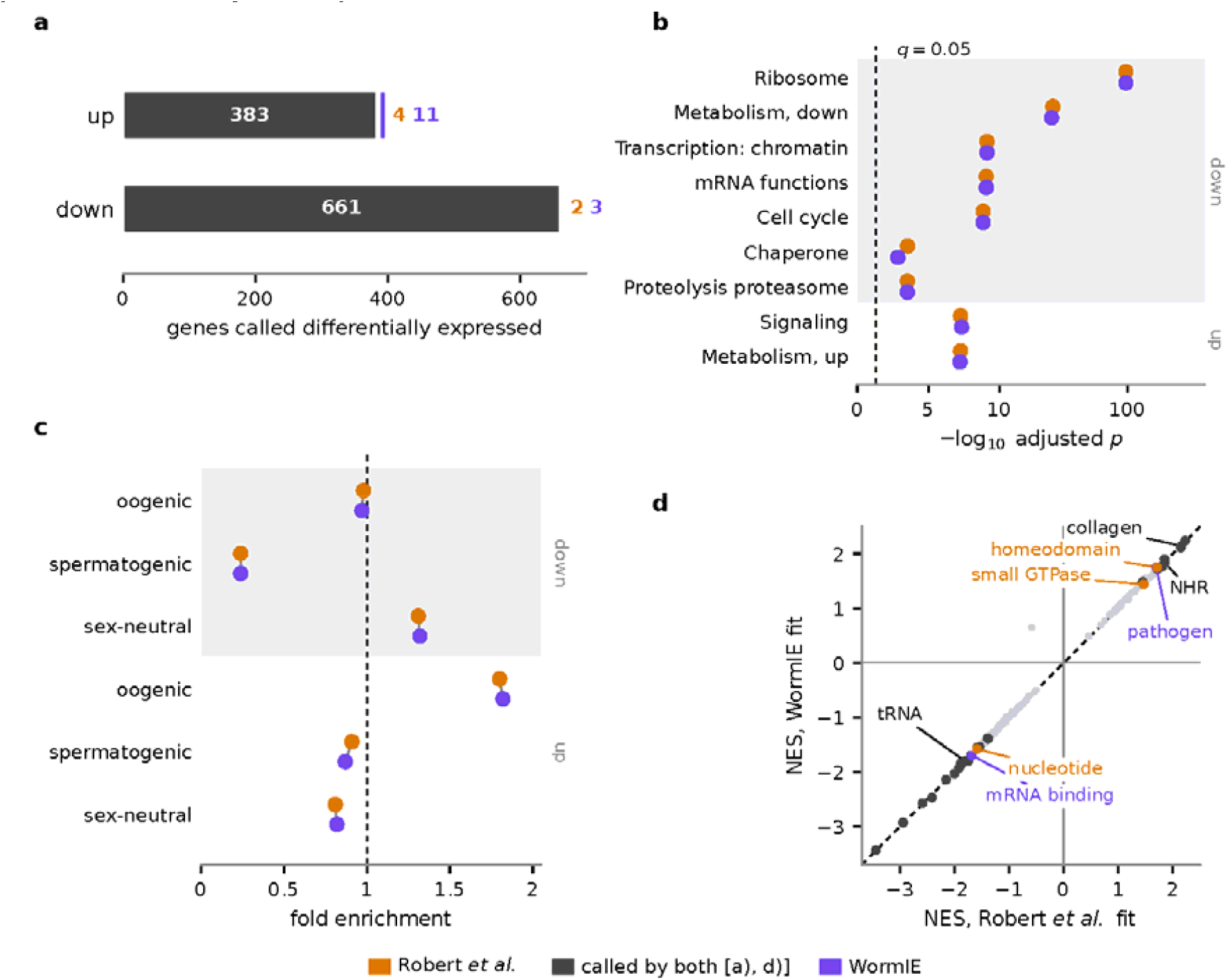
A published C. elegans germline RNA-seq analysis, repeated in WormIE. Wild type against sin-3(syb2172), from the raw reads of Robert et al. (2023), with that paper’s own gene selection applied throughout: nuclear protein-coding genes with a mean normalized count above 10, called differentially expressed at an adjusted p-value below 0.05. The key below the panels applies to all four: purple is WormIE and orange Robert et al. (2023). In b) and c), shaded bands mark the down- and up-regulated sets, labelled at the right, and the two dots in each row are drawn slightly apart so neither hides the other. a) Genes called differentially expressed, split by direction. Each bar is the union of the two analyses, divided into the genes both call in grey, the genes only WormIE calls in purple and the genes only Robert et al. (2023) call in orange. The white number counts the shared genes, and the colored numbers beside each bar count the genes only one analysis calls. The totals are therefore 394 against 387 up, and 664 against 663 down. No gene is called significant by both in opposite directions. b) Every WormCat category the published analysis reports, plotted as the adjusted p-value each run assigns it (hypergeometric test, Benjamini-Hochberg correction). Both runs recover all nine, and the axis is scaled so that the ribosome result does not compress the rest. c) The germline classification of Ortiz et al. (2014), which the published analysis reports in its Fig. 3D, plotted as fold enrichment rather than as significance. The test is one-sided, so a depletion cannot reach significance: spermatogenic genes are four times rarer among the down-regulated set than chance would give, at a fold of 0.24, where the significance axis of b) would show nothing. The two runs land within 0.04 of each other on all six sets. d) Ranked-list enrichment of all 168 WormCat categories tested, on the genes of each fit ranked by their Wald statistic, plotted as the normalized enrichment score (NES) each fit gives a category. Grey points are significant in both fits at an adjusted p-value of 0.05 or below, purple and orange in one fit only and pale points in neither. The scores fall on the diagonal (Pearson r = 0.995), and the 28 categories significant in both fits move in the same direction. Labels in black name the three categories the ranked test calls in both fits but over-representation calls in neither (Supplementary Note 11), and labels in purple or orange name the five called in one fit only, each close to the 0.05 cutoff in both.

The published analysis counted reads with the Python program htseq-count (Anders et al. 2015), whereas WormIE uses featureCounts (Liao et al. 2014). The model and the enrichment categories are otherwise the same: DESeq2 for the fit and WormCat’s whole-genome categories for the enrichment. Neither analysis shrinks fold changes, and both counted reads against the WS278 WormBase gene models. The strandedness of the libraries is not given in the paper, so WormIE counted unstranded, and the library totals then matched the authors’ deposited counts. Across the six libraries, gene-level counts from the two counters agree closely (r > 0.998), with library totals differing by less than 0.15%. The published expression filter, nuclear protein-coding genes with a mean normalized count above 10, retained nearly the same genes in both runs: 7,569 from WormIE’s counts and 7,555 from the authors’ results table, with 7,553 shared. Supplementary Note 11 gives both pipelines in full alongside the agreement at each step.

For differential gene expression, we fitted the authors’ model, genotype plus the day each replicate was prepared (as described in their published R script), and kept DESeq2’s defaults as that script does. WormIE identified 99% of the genes the published analysis called up and more than 99% of those it called down (Figure 3a), and no gene was called significant in opposite directions. Run on the authors’ own counts, WormIE recovered 386 of their 387 up-regulated and 662 of their 663 down-regulated genes and called no others. The authors’ model, run in native R with DESeq2 1.52.0, reproduces their published calls exactly, so the eight Bioconductor releases between their DESeq2 (1.36.0) and the one WormIE carries change nothing here. The one gene missed in each direction on the authors’ counts most likely reflects their removal of low-count genes after the fit, and the remaining differences in the full run would then come from the read counter.

The biological conclusion is unaffected. WormIE’s enrichment analysis identified every WormCat category reported by Robert et al. (2023), in the same directions, with seven categories among the down-regulated genes and two among the up-regulated genes (Figure 3b). Genes down in the mutant were most strongly enriched for ribosomal functions, at an adjusted p-value below 10^−97^, followed by metabolism, chromatin-associated transcription, mRNA functions, cell cycle, chaperone and proteolysis proteasome. Genes up in the mutant were enriched for metabolism and signaling.

Robert et al. (2023) also ran a second enrichment test, on the germline gene classification of Ortiz et al. (2014). WormIE supports this type of analysis through Custom gene set testing, which accepts user-supplied gene sets in place of its built-in collections. Among the genes upregulated in the mutant, WormIE’s version of the same analysis identified only the oogenic gene set as enriched, as in Robert et al. (2023), with the same fold enrichment of 1.8 as the published gene list (Figure 3c).

Beyond the two tests Robert et al. (2023) ran, WormIE can take the same data further. WormIE’s ranked-list enrichment follows the gene set enrichment analysis (GSEA) of Subramanian et al. (2005). It scores each category on the ranking of every measured gene rather than on the genes past a significance cutoff, so it can detect a category whose genes each shift too little to pass the cutoff alone. In WormIE this additional analysis takes just a few clicks, since one button on the differential expression results sends every gene, ranked by its Wald statistic, to ranked-list enrichment. We ran it on WormIE’s fit and on the authors’ results table, and the two gave nearly the same score to every WormCat category (Figure 3d). It recovered four of the seven categories reported among the down-regulated genes and signaling among the up-regulated ones, and it added three more significant categories (Supplementary Note 11).

In summary, repeating the analysis end to end reached the same biological conclusions as the published one, and the few genes on which the two runs disagree trace to the read counter and to when low counts are filtered.

## Discussion

WormIE (https://wormie.bio) is a gene-centric knowledgebase and analysis environment. It integrates seventeen annotation domains on a single gene identifier, so a query can begin with a gene, phenotype, tissue or interaction partner and retrieve related information without leaving the tool. A gene page gathers what those domains hold about one gene, including four panels to identify which other genes it works with: two drawn from recorded and predicted interactions and two computed by WormIE from annotation and expression data. Gene lists can be resolved, compared with one another and carried into over-representation or ranked-list enrichment across nine built-in collections. A collection supplied by the user can stand in for any of them and be tested in the same way. WormIE also runs the usual RNA-seq analysis route against that same annotation, from read counting through differential expression to enrichment.

Both browsing and analysis happen in a browser tab with no server behind them, so data provided by a user never leaves their machine. Some analyses run the standard package itself, compiled for the browser, and the rest are implemented with code written for WormIE that works directly against the local database. Analyses of simulated data in WormIE closely agreed with the same data analyzed in native R. Repeating a published germline analysis end to end recovered all nine of the WormCat categories reported in the original study and reached the same biological conclusion.

Most of the annotations in WormIE come from established community resources, chiefly WormBase (Davis et al. 2022) and the Alliance (Agapite et al. 2022), so the annotation behind an answer carries the authority of the projects that curate it. The application, database and build script are all released publicly, so a laboratory can rebuild WormIE from the same upstream sources or host the application independently, including on an internal network without internet access.

The public instance is maintained by the corresponding author using funding that covers the next several years, with the Richardson laboratory as a second, longer-term source of support. The price of that maintenance is small, since the recurring costs are simply a domain and the storage that holds the database for download, and every analysis runs on the user’s own machine rather than on hardware anyone has to keep funded. Should the hosted copy lapse, the build script, the application and the database remain public, and a laboratory that has already downloaded a build keeps working from it offline. We also plan to archive every published build, so that a result exported from any build of WormIE can be reproduced against the exact data behind it, long after the site has moved on to newer builds.

Most of the annotation WormIE carries describes the whole genome, where only two curated sets answer one narrow question each: the neuronal genes taken from WormBook and the synapse development and function classification curated for this work. Classifications of that kind are scattered through the literature and finding more of them is a matter of time rather than of method, so later builds should carry more than two. A user who assembles a collection for Custom gene set testing and finds it worth keeping is invited to send it, so that it reaches the community rather than staying on one machine. Corrections to the synaptic genes and suggestions for additional genes are welcome.

What the interface should do next is better answered by the people using it than by us. Once WormIE has been in use for a while we intend to ask the community what is awkward, what is missing and what a worm laboratory needs that no tool currently provides.

## Data availability

WormIE runs at https://wormie.bio in any current browser, with no installation, account or upload. Nothing a user analyses leaves their machine, so no data submitted to WormIE is held by the authors. The application source, the database build script and the curated gene sets distributed with it are at https://github.com/rayandrew/wormie.rs.ht under the MIT license.

Every source the database draws on is public, and Table 1 gives the release or retrieval date of each. The build reported throughout this manuscript is 20261025-112346, which reads the Gene Ontology release of 2026-07-26 and WormBase WS298, the final major release before that project moved to archival maintenance. Each build writes a version file recording the release of every source it read, and the methods text and citation list WormIE exports name the build, so any figure reported here can be traced to the data behind it. The site serves only the newest build, so the build reported here is archived at Zenodo under **[pending]**. An older build also survives in the browser that downloaded it, until the user clears the site’s data or an update to the application needs a newer database structure.

The exported methods text also names the build and the versions of the R packages behind a differential expression result (DESeq2, apeglm and ashr in Table 2), and WormIE keeps each set it has shipped, since the repositories distributing those packages serve only their current versions.

The harness that compares WormIE against the original packages in native R is in benchmark/ in the application repository, alongside the two reports this manuscript draws on, results/report.md and results/report-go.md. The complete result tree, including the per-term output behind those reports, is archived at Zenodo under **[pending]**. Supplementary Note 10 describes what was measured and how the two arms were matched.

The reanalysis in *A published analysis, reproduced in a browser* uses the germline RNA-seq of Robert et al. (2023), available from GEO under accession GSE227368, a subseries of GSE227499, and from BioProject PRJNA944713. The runbook, the derived count tables and the gene lists that section compares are archived at Zenodo under **[pending]**, and Supplementary Note 11 describes the comparison step by step.

The manuscript sources, including every figure script, are at **[pending]**.

## Acknowledgments

We thank WormBase for the gene, phenotype and literature curation it built up to its final major release, and the Alliance of Genome Resources, where that curation now continues. We also thank the Gene Ontology Consortium, CeNGEN, BioGRID, STRING, the WormCat authors and the other sources in Table 1 for making their data freely available. We thank Robert et al. (2023) for depositing their sequencing reads, results table and analysis script, which made the reanalysis in this paper possible. We also thank Ruiling Zhong and the other past and present members of the Richardson lab, who adopted WormIE early, tested its functions and gave the feedback that shaped it. We used Claude Fable 5.1 (claude-fable-5-1) and Claude Opus 5.5 (claude-opus-5-5), both from Anthropic, to refine the writing and the code. We also used them to scan the literature at the start of curating the annotation sets. The authors verified the curated assignments manually and take responsibility for all content.

