## Supplemental figures and notes for "WormIE: an Integrated Explorer for multi-source *Caenorhabditis elegans* annotations and transcriptomic data interpretation"

### Supplementary Figures

The screenshots in Supplementary Figures S1 to S8 were taken from the public site in Chrome, on database build 20261101-012552, with a window 800 pixels wide, so the application lays itself out as it does on a narrow screen and its text stays legible at print size. Supplementary Figure S6 used a window 1,280 pixels wide, so that its plots sit two to a row. They were captured at four times the screen resolution and are printed at 300 dots per inch. Each is cropped to the part of the page its caption describes, and anything else added or hidden for a figure is stated in its caption.

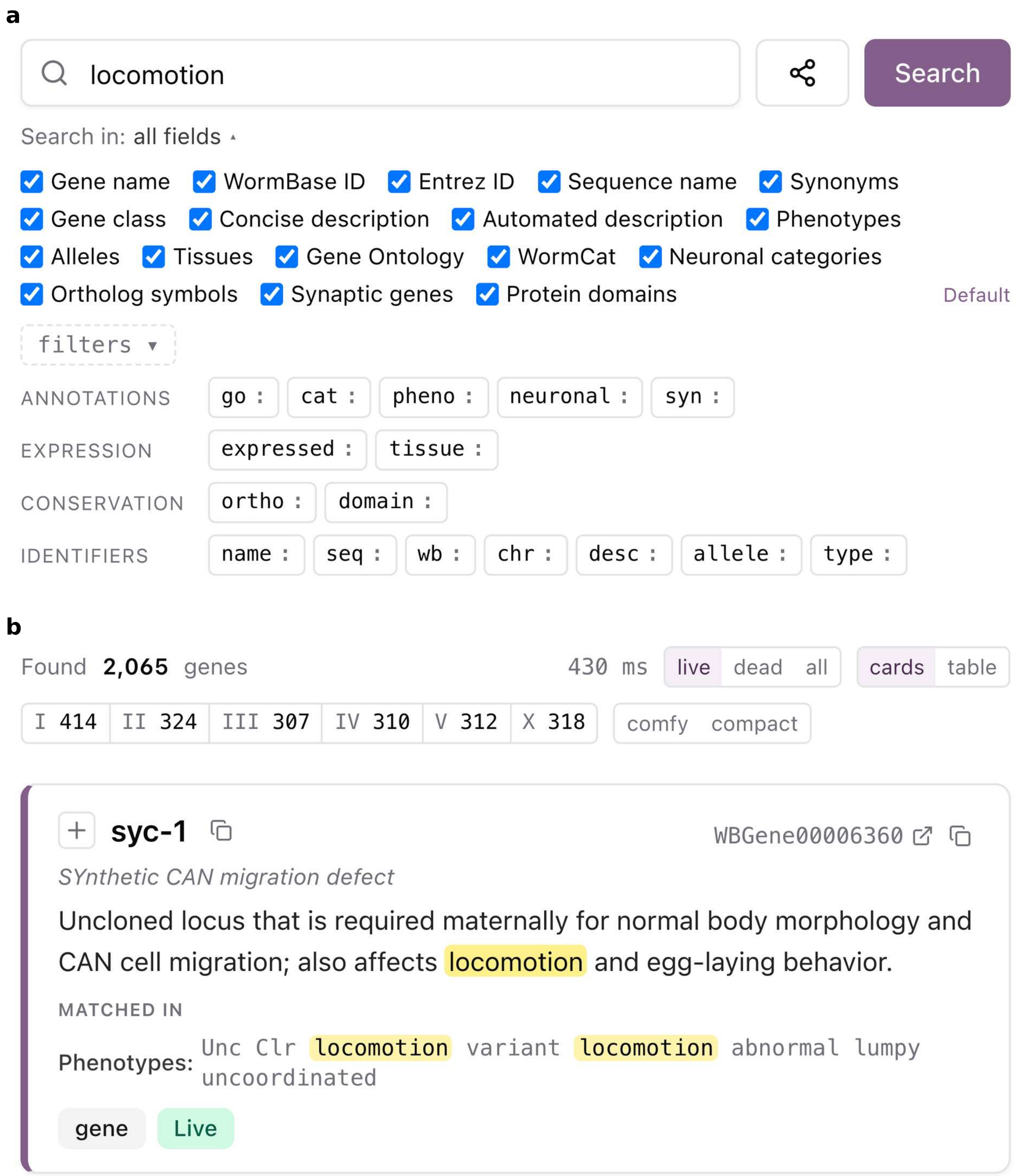


Supplementary Figure S1: **The whole database behind one search box.** With every field ticked under Search in, one query searches gene names, identifiers, descriptions, phenotypes, alleles, tissues, GO terms, WormCat categories, neuronal genes, synaptic genes, protein domains and ortholog symbols. a) The search for `locomotion`, with the filter prefixes open, `chr` for chromosomes among them. b) Facets for gene status and chromosome narrow the 2,065 genes found, and the first matches in both its description and its phenotypes. The buttons that send results to an analysis are cut from between the two parts.

Alt text: Two screenshots of the search page. a) A search box holding the word locomotion, a grid of ticked boxes naming the fields searched and rows of filter prefixes. b) The result count, buttons that filter by gene status and chromosome and the first matching gene with its description and phenotypes highlighted.

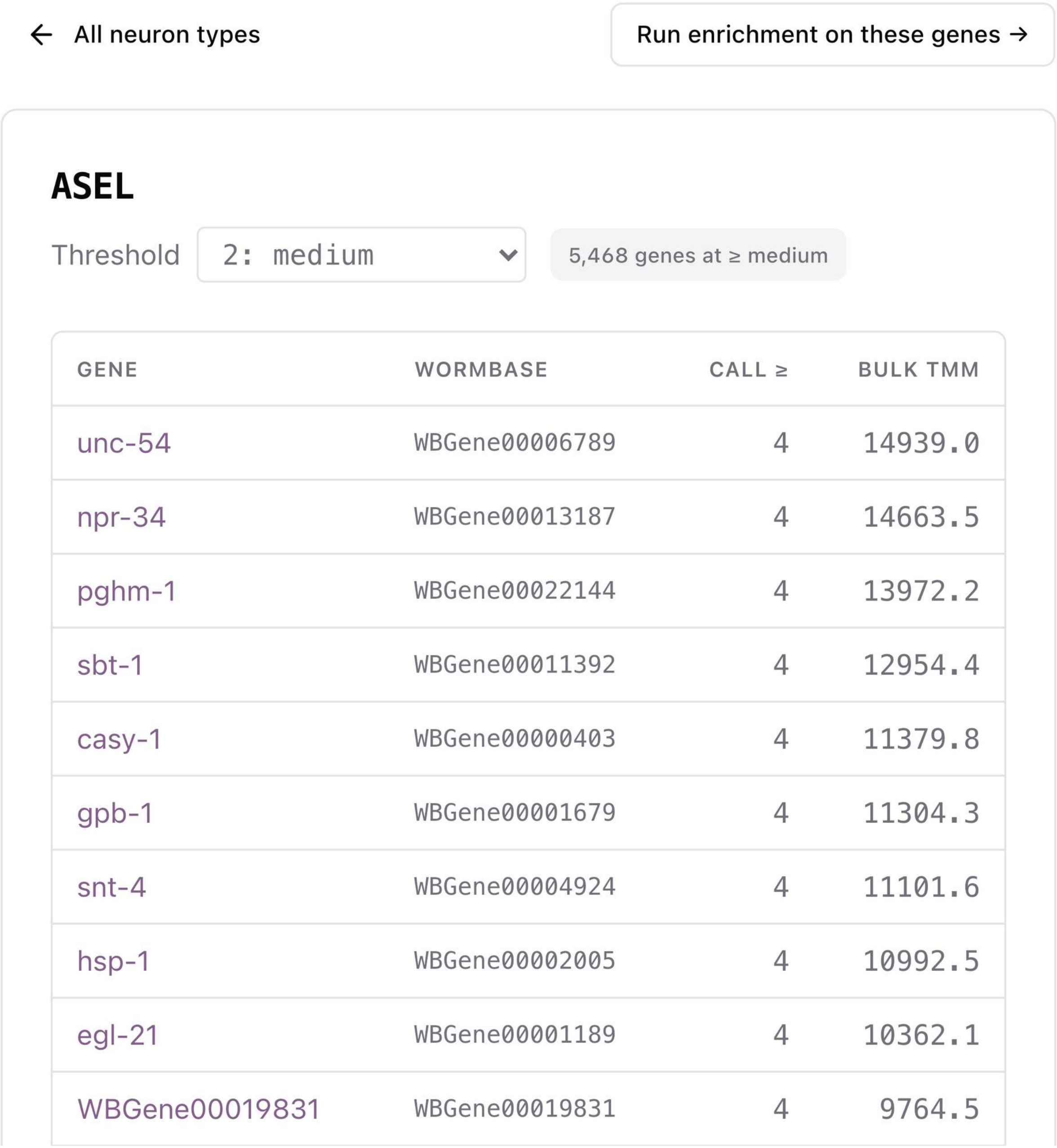


| GENE | WORMBASE | CALL ≥ | BULK TMM |
|---|---|---|---|
| unc-54 | WBGene00006789 | 4 | 14939.0 |
| npr-34 | WBGene00013187 | 4 | 14663.5 |
| pghm-1 | WBGene00022144 | 4 | 13972.2 |
| sbt-1 | WBGene00011392 | 4 | 12954.4 |
| casy-1 | WBGene00000403 | 4 | 11379.8 |
| gpb-1 | WBGene00001679 | 4 | 11304.3 |
| snt-4 | WBGene00004924 | 4 | 11101.6 |
| hsp-1 | WBGene00002005 | 4 | 10992.5 |
| egl-21 | WBGene00001189 | 4 | 10362.1 |
| WBGene00019831 | WBGene00019831 | 4 | 9764.5 |

Supplementary Figure S2: **An annotation page as a starting point.** Every GO term, phenotype, tissue, neuron type and human disease has a page of its own listing the genes annotated to it, so a search can begin at a term rather than at a gene. Shown is the page for the ASEL neuron type, which lists the 5,468 genes CeNGEN calls expressed there at its medium threshold, ranked by expression in sorted bulk samples, with a button that sends them to enrichment analysis. The help text between the button and the list is cut out.

Alt text: A screenshot of the page for the ASEL neuron type, with a threshold menu and a table of genes, their WormBase identifiers and their expression values, and a button that runs enrichment on the genes.

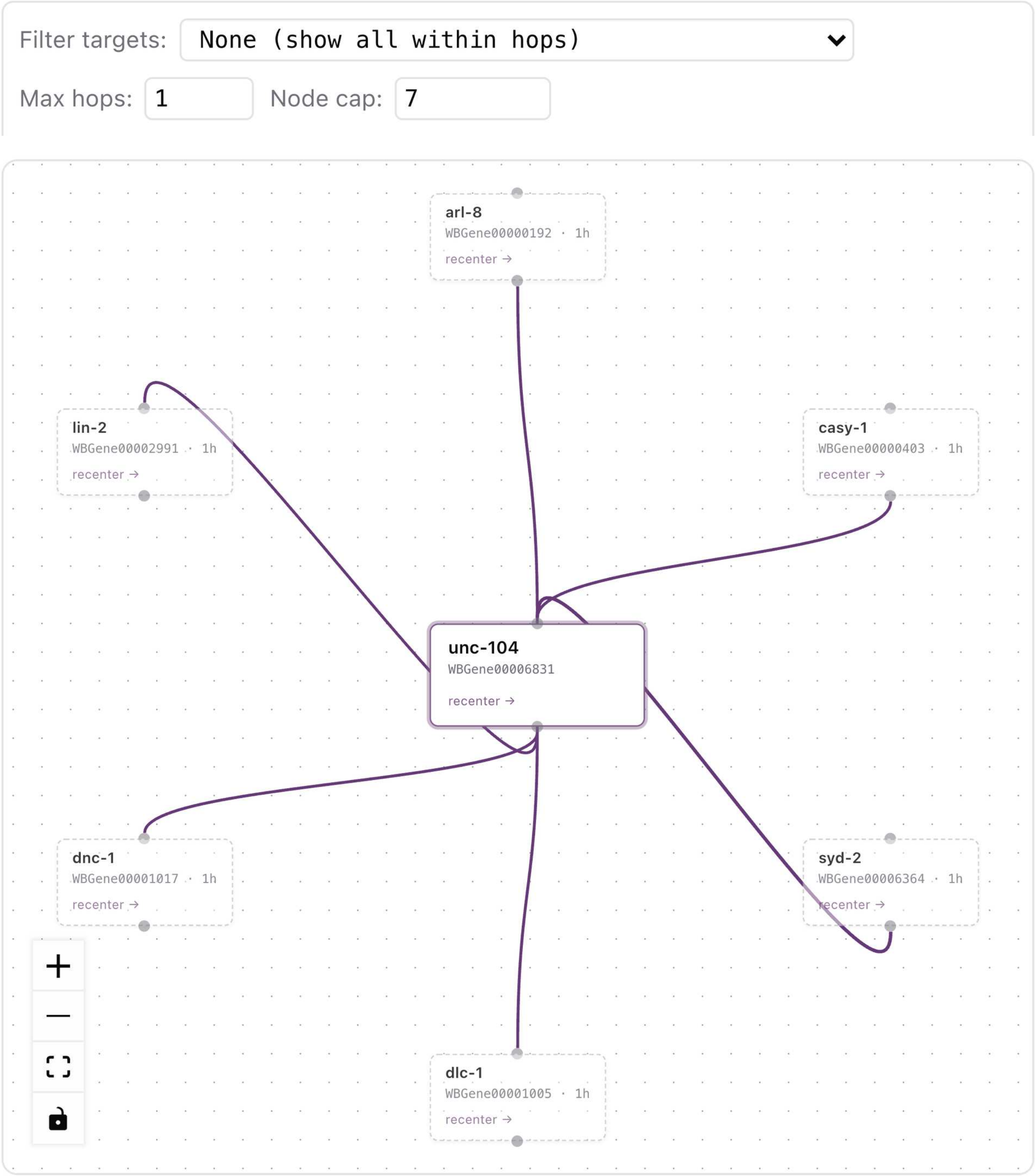


Supplementary Figure S3: **Interaction partners as a navigable neighborhood.** The experimentally recorded interactions from BioGRID open as a graph, which can be explored outward from any node rather than read as a list. Shown are *unc-104* and six of its direct partners, and *recenter* on any node makes that gene the center. The controls above the graph widen the neighborhood to partners of partners (Max hops) and restrict it to a neuronal gene category, a WormCat category or a saved list (Filter targets). The node cap is set to seven here so that the gene names stay legible, and the overview map is hidden.

Alt text: A screenshot of a network graph with unc-104 in a box at the center and six partner genes in boxes around it, joined to it by purple curved lines, under two boxes that set the number of hops and the node cap.

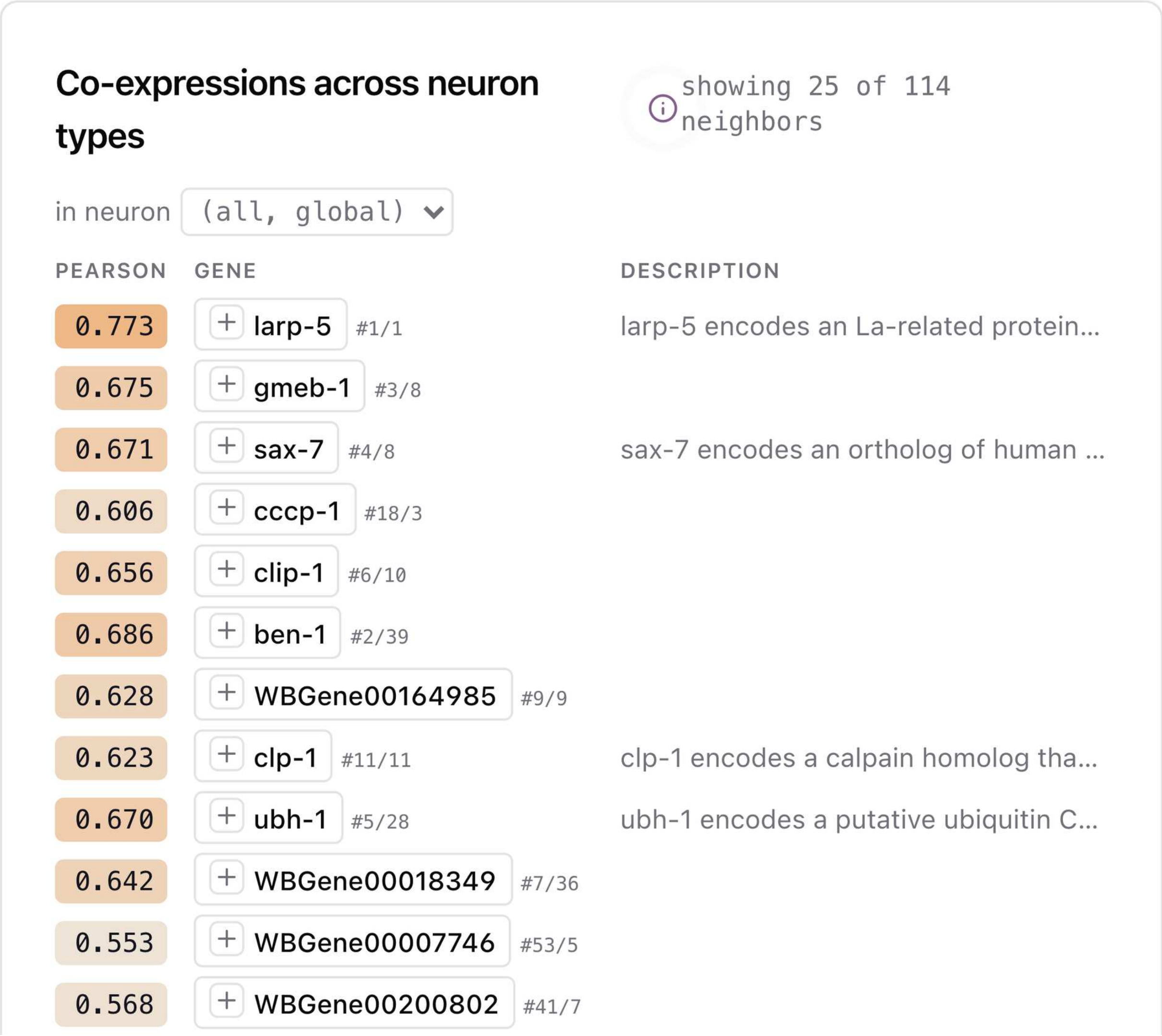


Supplementary Figure S4: **Neighbors ranked by co-expression rather than by annotation.** The expression profile of the query gene is set against that of every other gene across the 52 neuron types CeNGEN covers, so two genes can rank as neighbors without sharing a single annotation. Shown are the first 12 of the 114 neighbors of *unc-104*, each with its Pearson correlation. The two numbers after each name are the neighbor's rank in the list of *unc-104* and the rank of *unc-104* in the neighbor's list. For example, #4/8 after *sax-7* means that *sax-7* is the fourth neighbor of *unc-104* and that *unc-104* is the eighth neighbor of *sax-7*, and #1/1 marks a pair that each ranks first (Supplementary Note 6). Several of the neighbors are uncharacterized genes.

Alt text: A screenshot of a table of neighbors of unc-104 with a Pearson correlation in an orange box, a gene name and a short description in each row, with the strongest correlations at the top.

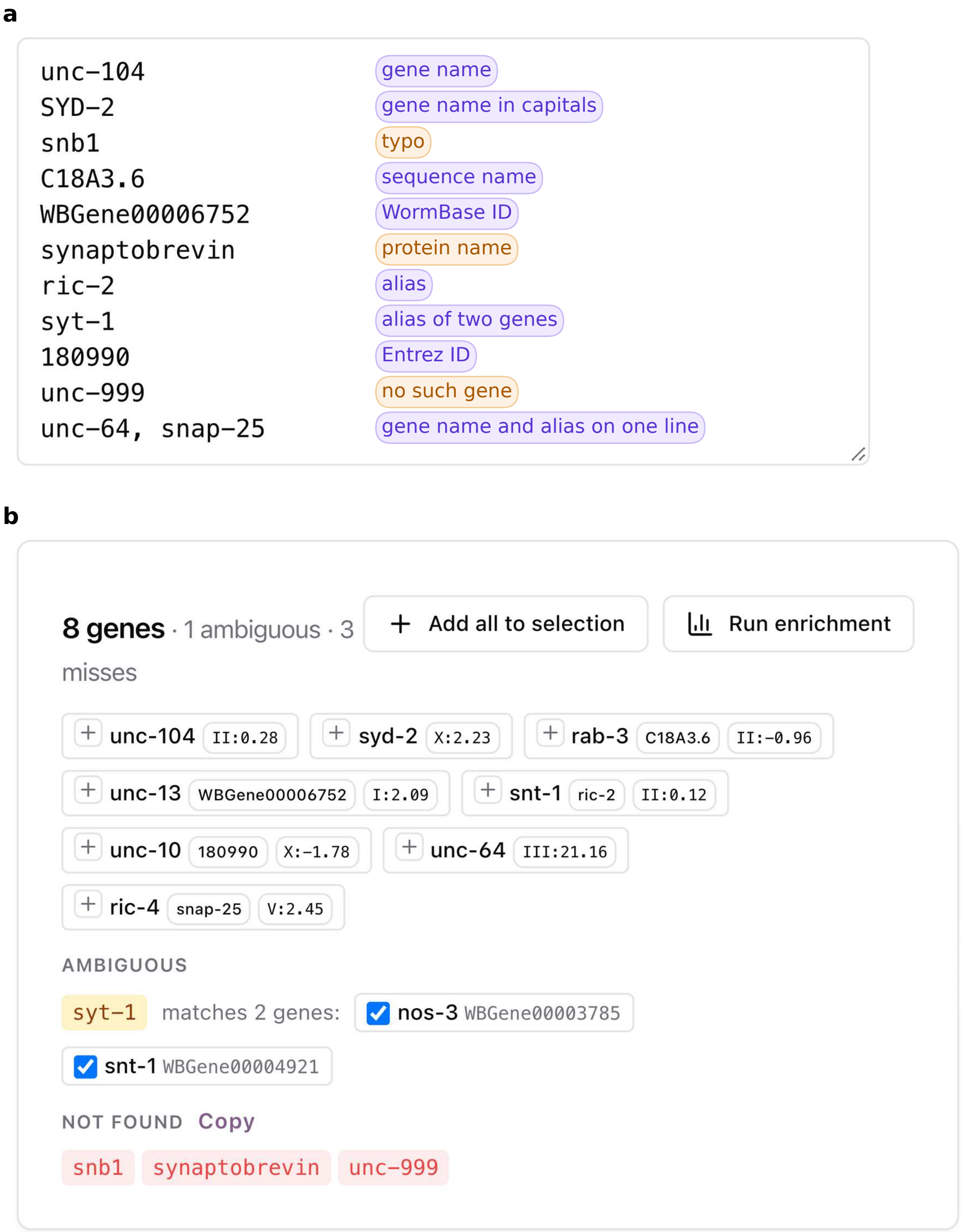


Supplementary Figure S5: **Mixed identifiers resolved in bulk.** Gene names, sequence names, aliases, WormBase identifiers and Entrez IDs pasted together are each mapped to a WormBase identifier. a) A made-up list from a synapse screen. The colored labels beside each entry were added to the screenshot for this figure, and WormIE does not show them. The three entries labelled in orange name no gene. b) The result as WormIE shows it, where each match shows the entry that produced it. The alias *syt-1*, which WormBase gives to both *snt-1* and *nos-3*, is listed apart as ambiguous with both genes ticked, so the user can untick the one not meant.

Alt text: Two screenshots of the identifier tool. a) A box holding ten pasted entries, each with a colored label that names its kind, added to the screenshot for this figure. b) The result, showing eight genes found, one ambiguous alias with two ticked options and three entries not found in red.

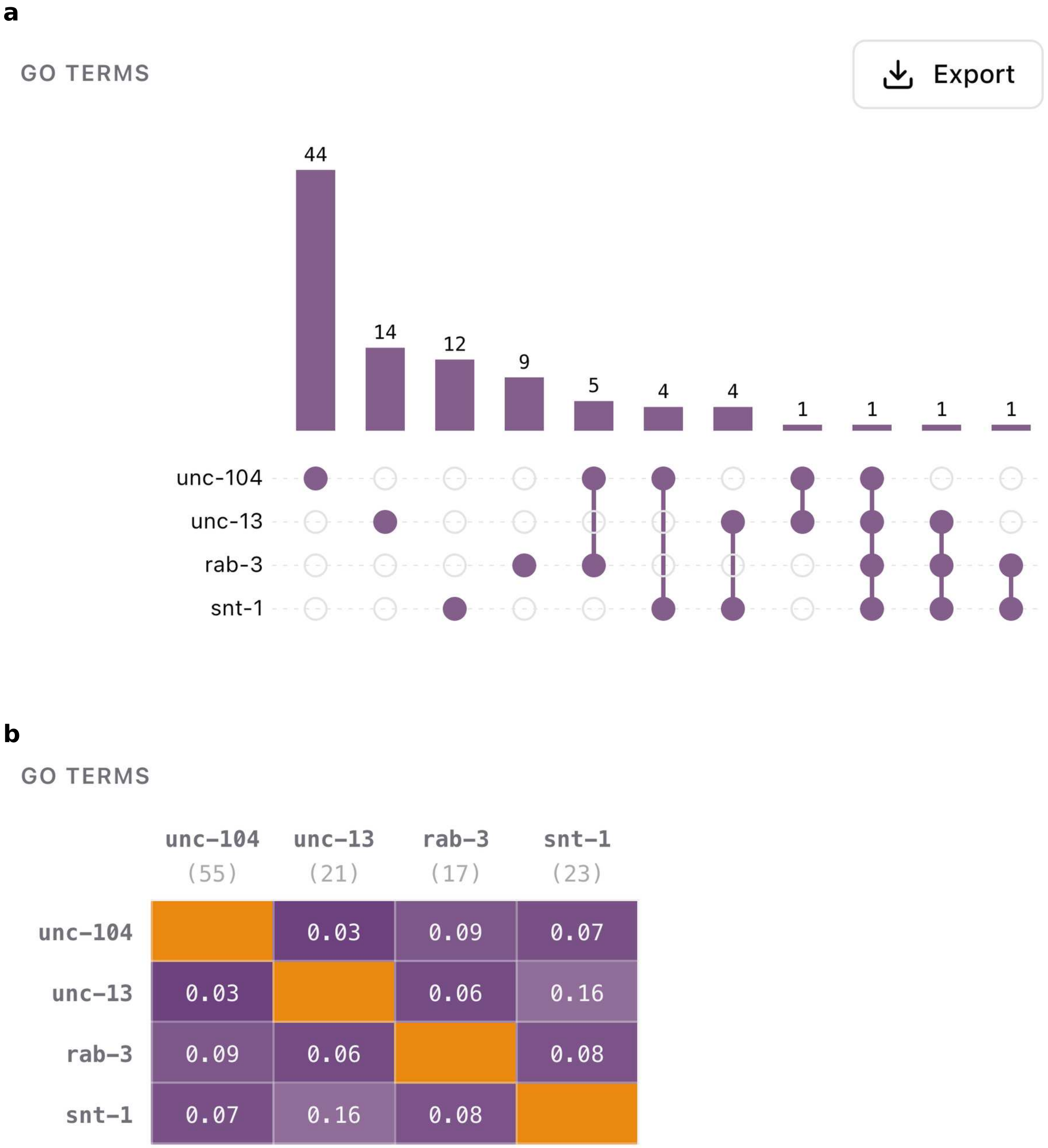


Supplementary Figure S6: **Several genes compared on the annotations they share.** The Annotation overlap page for four genes that act at the synapse, *unc-104*, *unc-13*, *rab-3* and *snt-1*. For three or more genes the page draws an UpSet plot and a Jaccard heatmap for each annotation type, of which the Gene Ontology pair is shown here, and lists the annotations common to every gene. a) The GO-term UpSet plot. Each bar counts the terms held by exactly the genes whose dots are filled below it, so 44 terms belong to *unc-104* alone. b) The GO-term Jaccard heatmap, where each cell is the share of their terms that two genes hold in common and the number under each gene is its count of terms. No pair shares more than 0.16 of its terms, and the only GO term common to all four is plasma membrane. Beyond GO, the page lists one phenotype (locomotion variant), three tissues and 120 CeNGEN neuron types common to all four, and no shared ortholog. All four are detected in nearly every neuron type yet share a single GO term.

Alt text: Two panels for four genes. a) An UpSet plot, with a bar chart of term counts above a grid of filled dots that show which genes share them. b) A square heatmap of the share of terms two genes hold in common, with low values for every pair of different genes.

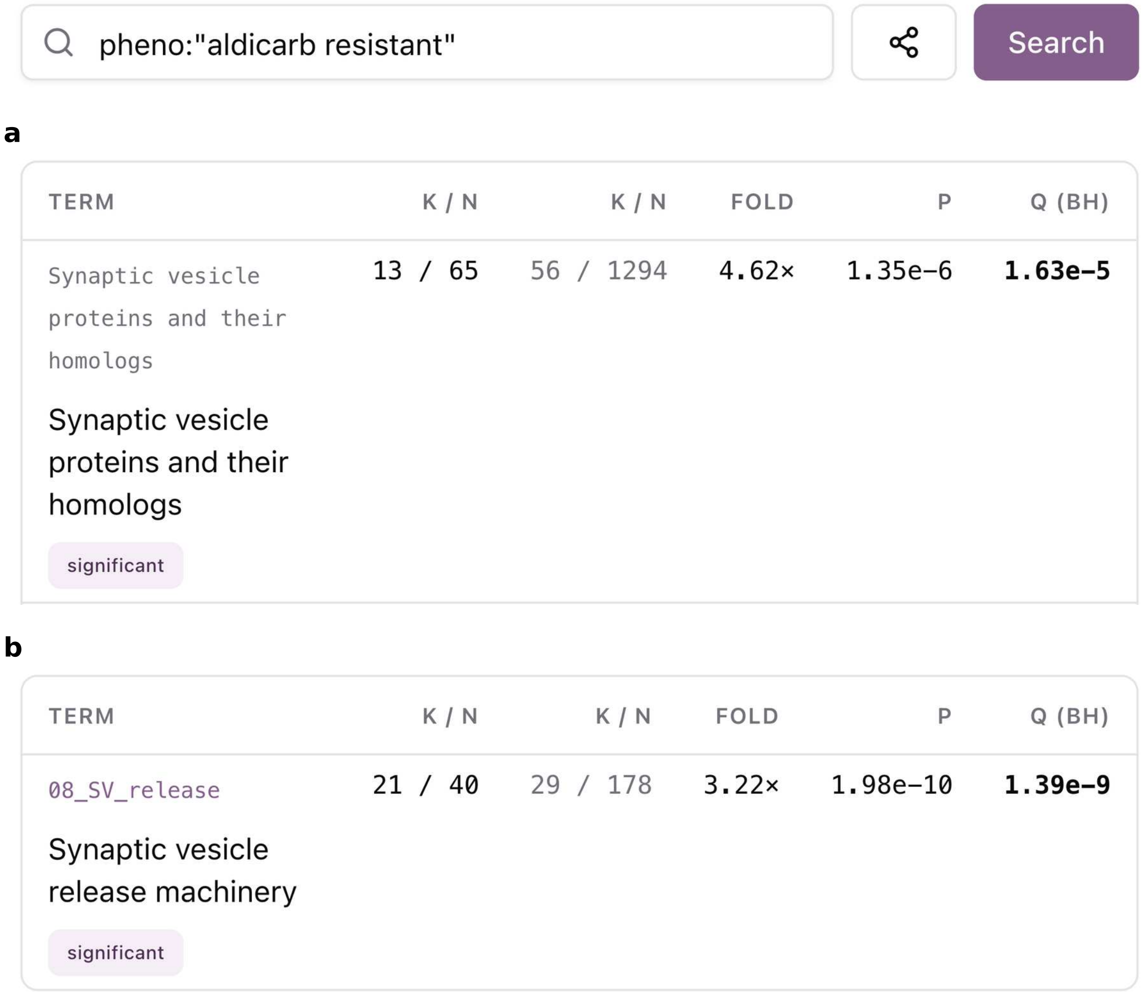


Supplementary Figure S7: **One result from each of the two collections found in no other worm resource.** The gene list is what a phenotype search returns, sent on with the Over-representation button under the search results. The search box is shown once, above both results. The 250 genes found by `pheno:"aldicarb resistant"` are tested against the neuronal genes (a) and the synaptic genes (b). a) The neuronal genes cover 65 of the 250 genes, and 3 of the 12 categories tested are significant. The category of synaptic vesicle proteins and their homologs leads, with 13 of those 65 genes in a category of 56, a 4.6-fold enrichment at an adjusted p-value of $1.6 \times 10^{-5}$. b) The synaptic genes cover 40 of the 250 genes, and the synaptic vesicle release machinery class is the only significant one of seven tested, holding 21 of those 40 genes, a 3.2-fold enrichment at an adjusted p-value of $1.4 \times 10^{-9}$.

Alt text: Two screenshots of enrichment results under a phenotype search box. a) One row for the synaptic vesicle protein category with its counts, fold, p-value and adjusted p-value. b) One row for the synaptic vesicle release machinery class with the same columns.

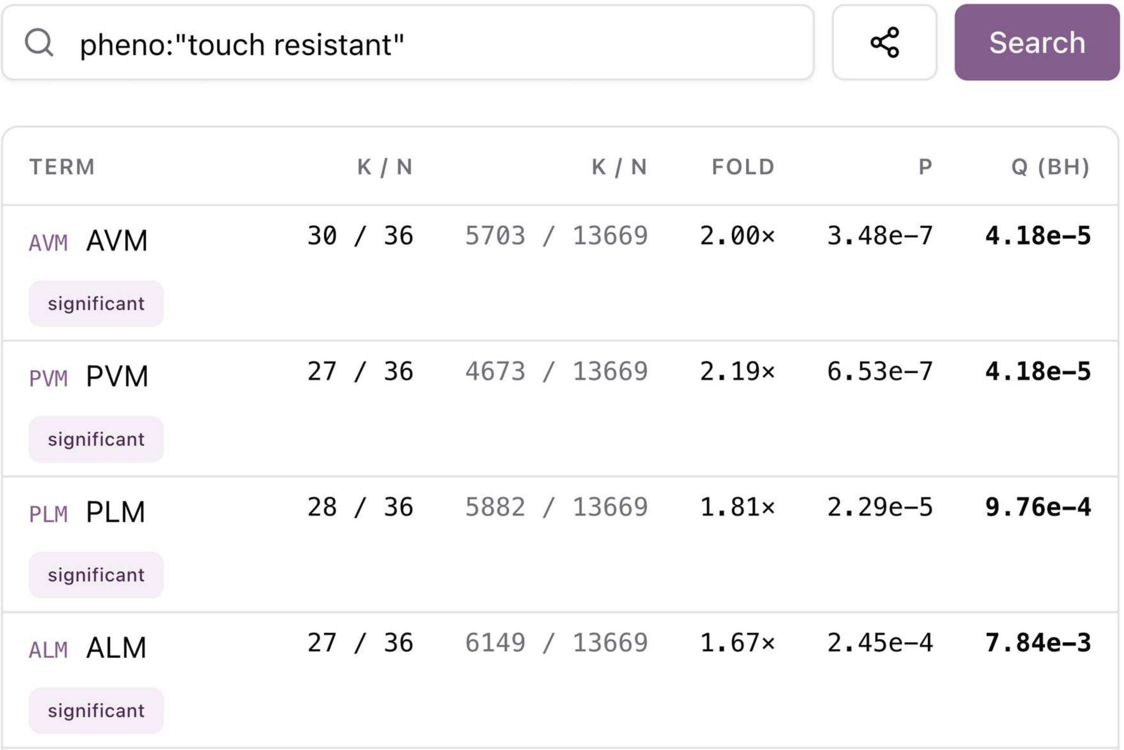


| TERM | K / N | K / N | FOLD | P | Q (BH) |
|---|---|---|---|---|---|
| AVM AVM significant | 30 / 36 | 5703 / 13669 | 2.00× | 3.48e-7 | **4.18e-5** |
| PVM PVM significant | 27 / 36 | 4673 / 13669 | 2.19× | 6.53e-7 | **4.18e-5** |
| PLM PLM significant | 28 / 36 | 5882 / 13669 | 1.81× | 2.29e-5 | **9.76e-4** |
| ALM ALM significant | 27 / 36 | 6149 / 13669 | 1.67× | 2.45e-4 | **7.84e-3** |

Supplementary Figure S8: **One result from the CeNGEN neuron types.** The 42 genes found by `pheno:"touch resistant"` are sent on with the Over-representation button under the search results and tested against the CeNGEN neuron types. The classes cover 36 of the 42 genes. The four touch receptor neuron types, AVM, PVM, PLM and ALM, lead the seven significant types of 128 tested, and ALN, PVD (a harsh-touch neuron) and IL1 follow below the cut.

Alt text: A screenshot of a results table with the neuron types AVM, PVM, PLM and ALM in rows, each with its counts, fold enrichment and adjusted p-value.

### Supplementary Note 1. Implementation note

The main text describes what WormIE does for the reader without saying how. This note records how: where the computation runs, what the browser stores, how several tabs share one database and how the site and its database are built and served. A user does not need this information to use WormIE or to interpret its output.

#### Where the computation happens

WormIE is a set of files served as they are: there is no server program running behind the site. Everything runs inside the browser tab on the user's own processor. A browser tab normally handles one task at a time, and responding to the user is one of those tasks, so a long calculation leaves the page unable to respond until the calculation ends. WormIE instead hands every calculation to a worker, a second task runner the browser provides for exactly this purpose, so the page keeps responding and shows results as they arrive.

The database is SQLite, which reads only the parts of the file a query touches, so a 598 MB database can be searched efficiently in a browser tab. That capability is what makes the lookup-based analyses (over-representation, ranked-list enrichment and per-sample gene set scoring) straightforward to rewrite for the browser (Supplementary Note 10). A feature that needs more than search, such as DESeq2, runs the original package instead. This is possible through WebAssembly, a format that lets a browser execute programs that would otherwise have to be installed on the computer. R and DESeq2 are compiled to WebAssembly from their unchanged source code. The browser therefore runs the same R and the same packages as a laptop, inside a worker like everything else. Supplementary Tables S6 and S7 give the trade-offs of running R this way.

#### Storage and working offline

On the first visit the browser downloads the database and keeps it on the computer for SQLite to read. R and the program that counts reads are fetched the first time an analysis needs them, about 100 MB for R and its packages (Supplementary Table S7), and stay on the computer as well. The pages of the application are kept the same way, so a later visit reads everything from the machine and can work offline. A connection is needed only to fetch a newer database build or a newer version of the application, and WormIE asks before switching to either. A user's saved information, such as notes, gene lists and bookmarks, is preserved on the computer as well. Nothing a user enters leaves the computer, because WormIE has no user accounts and no server to receive the data.

WormIE asks the browser to treat its storage as permanent by default. Even so, a browser running low on disk space may delete the stored files, and the next visit then downloads the database again with the saved notes and gene lists cleared. Two habits guard against this. One is to install WormIE as an application: Chromium browsers count installation among their grounds for granting permanent storage without asking (LePage 2020), and Safari gives an installed web app a larger storage quota. The other is to export the marked genes and their notes as a spreadsheet at the end of each session; the identifiers can be pasted into the *Gene name resolver* to rebuild the list, though the notes have to be re-entered by hand.

#### Several tabs at once

A user often has several WormIE tabs open, for example a gene page beside a running analysis, and opening the database in every tab would cost its memory several times over. WormIE avoids this by letting only one tab hold the database open and having the other tabs send their queries to it. If that tab closes or stops responding, the others notice and one of them takes over.

#### Rebuilding the database

One script rebuilds the database automatically from public sources, weekly and whenever the build script or the curated files change. It downloads every source itself, except three files kept with the application code: the WormCat whole-genome assignments, the neuronal gene catalog and the synapse development and function classification of Supplementary Table S1. Every source is downloaded afresh at each weekly rebuild, WormBase included, although its files no longer change since WS298 in November 2025 was its final major release. Alongside the compressed database, the script writes a record of the release or date of every source, the build date and the size of the database compressed and uncompressed.

The build reported in this manuscript is 598 MB uncompressed, and the browser downloads it compressed with Zstandard, at 83 MB. A 128 MB gzip copy of the same file is served beside it.

#### Hosting a copy of WormIE

A lab can host its own copy of WormIE on any ordinary web server. The server must send two settings with every file, `Cross-Origin-Opener-Policy: same-origin` and `Cross-Origin-Embedder-Policy: require-corp`, without which WormIE cannot open its database and may show no error. The public site adds a small helper program only because its hosting provider limits single files to less than the size of the compressed database.

The synapse development and function classification curated for this work is listed in full in Supplementary Table S1. Its twelve classes are ordered from synapse formation to function: the transcription factors that drive synaptic gene expression and the export of their transcripts, the cues that place a synapse along the axon and match it to its partner, active zone assembly and its growth limit, cargo transport, vesicle release, postsynaptic reception, neurotransmitter identity, electrical synapses and finally remodeling. Every gene cites at least one primary research article, shown beside the gene in the interface, and the reviews each class was assembled from are listed with the class. Each assignment was checked against the cited publications, but the classification is not exhaustive.

Supplementary Table S1: **The curated synapse development and function panel.** Each class, the number of genes assigned to it in the build reported in Table 1, the roles its genes are divided into and the publications its genes cite together with the reviews the class was assembled from. Reference numbers key to the list below.

| Class | Genes | Description | References |
|---|---|---|---|
| Activity-regulated transcription factors | 14 | Transcription factors, with their co-activators and homeobox cofactors, that promote the transcription of synaptic genes in response to neuronal activity. Roles: Co-activators and chromatin remodelers; Activity-induced factors; Homeobox cofactors. | 1, 5, 48, 49 |
| RNA export and processing | 6 | Genes encoding proteins involved in the export and processing of synaptic transcripts. Roles: THO complex subunits; Export adaptors and cofactors. | 5 |
| Synapse positioning | 7 | Genes encoding components of the Wnt pathway (ligands, receptors and transducers) that restrict synapse placement to the correct region along the axon. Roles: Wnt ligands; Wnt receptors and co-receptors; Intracellular transducers. | 8, 19, 20, 21 |
| Synaptic specificity and cell adhesion | 23 | Genes encoding proteins that direct synapses to the correct partner cell and hold them there. Roles: Adhesion and organizer molecules; Guidance cues and receptors; Cytoskeletal and signaling effectors. | 3, 13, 19, 31, 32, 33, 34, 35, 50, 51, 52, 53, 54, 55, 56, 57, 58, 59, 60, 61, 62, 63, 64 |
| Active zone scaffolding | 12 | Genes encoding the core active zone scaffolds and the kinases and phosphatases that act on them. Roles: Assembly regulators; Core scaffold proteins. | 3, 4, 11, 14, 18, 65, 66, 67, 68, 69, 70, 71, 72, 73 |
| Synapse growth control | 7 | Genes that limit synapse formation and terminate axon growth. Roles: PHR ligase complex and effectors; MAPK cascade kinases. | 3, 9, 10, 17, 46 |

| Class | Genes | Description | References |
|---|---|---|---|
| Synaptic cargo transport | 9 | Kinesin motors and the adaptors and regulators that carry synaptic vesicle precursors, dense-core vesicles and receptors along the axon. Roles: Kinesin motors; Motor adaptors and regulators. | 15, 74, 75, 76, 77, 78, 79, 80 |
| Synaptic vesicle release machinery | 29 | Genes encoding the fusion, release, recycling and acidification machinery of synaptic and dense-core vesicles, with the ion channels on either side of the synapse that gate release and excitation. Roles: SNARE and fusion machinery; Vesicle-associated proteins; Endocytosis and recycling factors; Neuropeptide processing enzymes; Synaptic ion channels; Vesicle biogenesis factors. | 3, 12, 16, 41, 42, 43, 44, 47, 81, 82, 83, 84, 85, 86, 87, 88, 89, 90, 91, 92, 93, 94, 95, 96, 97, 98, 99, 100, 101, 102 |
| Postsynaptic receptors and clustering | 32 | Genes encoding postsynaptic receptors of every receptor class, with the enzymes and scaffolds that cluster the receptors at their correct positions. Roles: Ionotropic receptors; Metabotropic receptors; Signaling and cleft enzymes; Scaffolds and clustering factors. | 2, 6, 12, 22, 23, 24, 25, 26, 27, 28, 29, 30, 103, 104, 105, 106, 107, 108, 109, 110, 111, 112, 113, 114, 115, 116, 117, 118, 119, 120, 121, 122 |
| Neurotransmitter identity | 14 | Genes encoding the components that give a neuron its neurotransmitter identity (cholinergic, GABAergic and so on). Roles: Vesicular transporters and cofactors; Biosynthetic enzymes; Reuptake transporters; Neuropeptide ligands. | 6, 12, 36, 37, 38, 39, 40, 123, 124, 125, 126, 127, 128, 129, 130 |
| Electrical synapses and innexins | 15 | Genes encoding the 15 innexins shown to be expressed in the nervous system and to rewire under stress and in dauer. Roles: Neuronally expressed innexins. | 7, 131, 132 |
| Remodeling transcription factors and effectors | 10 | Transcription factors, heterochronic timers and their effectors that set whether and when a neuron rewires its synapses. Roles: Pro-remodeling factors; Anti-remodeling factors; Heterochronic timers. | 2, 45, 133, 134, 135, 136, 137, 138, 139 |

### Supplementary Note 2. Gene Ontology propagation

The annotation file WormBase distributes records only the direct, most specific Gene Ontology term for each gene. During the build, WormIE propagates each annotation to the broader terms above it in the ontology, so that a gene annotated to a specific term also counts toward the broader terms. It follows the same rules as the `GOALL` mapping of GO.db (Carlson 2025) and the `enrichGO` and `gseGO` functions of clusterProfiler (Yu *et al.* 2012), so an enrichment result computed in WormIE is comparable with one computed in R from the same gene list and background:

- `is_a` and `part_of` propagate, since a gene acting in a specific process acts in the general one it is a kind of or a part of;
- `regulates`, `positively_regulates` and `negatively_regulates` propagate, since a regulator of a process is counted as involved in it;
- `occurs_in` and `has_part` do not, since neither implies that a gene annotated to the narrower term takes part in the broader one;
- an annotation with a NOT qualifier is left out of the propagated hierarchy and of enrichment backgrounds, since a NOT annotation rules out the term and its narrower terms but says nothing about its broader ones, and the annotation stays visible on the gene page.

Over-representation tests the three aspects of the ontology (biological process, molecular function and cellular component) separately. Pooling them would put a gene with no biological-process annotation into the universe of every biological-process test. That gene could never appear in any biological-process term, so it only makes the universe larger and every test more conservative. Each aspect therefore builds its own universe from the genes with at least one positive annotation in that aspect and runs its own Benjamini-Hochberg correction, as clusterProfiler, topGO and GOstats do. Ranked-list enrichment, whose universe is the ranked list itself whatever the aspect, corrects across all three at once, as fgsea does.

As a check on the propagation, we compared GO enrichment results from WormIE with those from clusterProfiler (Yu *et al.* 2012) using the class I targets of Tepper *et al.* (2013): genes up-regulated in a *daf-2* mutant, which map to 1,664 WormBase gene identifiers. WormIE reads a Gene Ontology release from July 2026 while the Bioconductor annotation package carries an earlier snapshot. Given the same gene list, WormIE recovers 149 of the 171 terms clusterProfiler calls enriched and identifies 35 additional terms. Seven of the 22 terms WormIE does not recover have been retired by the newer Gene Ontology release; for the other 15, the genes assigned to each term differ between releases. Of the 35 GO terms identified exclusively by WormIE, 31 differ in membership in the same manner. The remaining four carry the same genes in both tools, with adjusted p-values that fall just below the 0.05 threshold in WormIE and above it in clusterProfiler. Differences between annotation releases thus appear to explain most of the discrepancies. This comparison is one reason each build records the release date of every source it uses.

### Supplementary Note 3. Exclusions and filters at ingestion

Not every record in a source file enters the database. This note lists, source by source, what the build leaves out or thresholds and why, so that a count in Table 1 of the main text can be traced back to the file it was read from.

**NOT qualifiers.** Phenotype and anatomy annotations carrying a NOT qualifier are skipped as the database is built; the interface never displays them, so nothing visible is lost. Meanwhile, Gene Ontology annotations carrying that qualifier are kept, because the gene page does show them, and are treated as described in Supplementary Note 2.

**STRING.** STRING scores each predicted association from 0 to 1,000 by how much evidence supports it, and WormIE keeps only those scoring at least 700, the cutoff STRING itself labels high confidence.

**CeNGEN.** CeNGEN publishes a yes-or-no call for whether each gene is expressed in each of 128 single-cell neuron types at four thresholds, each one stricter than the last (Taylor *et al.* 2021). The database keeps the highest threshold each gene passes in each type. *daf-2*, for instance, passes thresholds 1 and 2 in the AWB neurons but not 3 and 4, and only threshold 1 in the SIB neurons. Neuron-type enrichment takes the genes detected at threshold 2 or above, the level CeNGEN uses throughout its own analyses, so *daf-2* belongs to the AWB gene set but not to the SIB one.

Separate from those calls, the expression values shown on the gene page and used for co-expression come from another CeNGEN source, the integrated matrix of July 2024 (Barrett *et al.* 2026). It covers 52 neuron types sequenced in bulk, combines those samples with the single-cell atlas and removes non-neuronal contamination from them. The single-cell atlas does not separate the DD and VD motor neurons, so the matrix, and therefore WormIE, reports them together as VD_DD. Contamination removal leaves a gene that is absent from a neuron type near zero rather than at zero, so values below 0.1 are treated as not detected and are not stored. CeNGEN's website identifies 17 genes that were overexpressed as transgenes in the sorted strains, and its 2024 analysis code filters a further 63 whose promoters drive the reporters used for sorting. The two lists overlap, so together they name 75 genes whose values in this matrix can be inflated. WormIE stores the values of these 75 genes and marks each one on its gene page. Co-expression pairs are not read from a file but computed by the build from those values; the rules that decide which pairs are stored are given in Supplementary Note 6.

**Publications.** Gene-to-publication links come from NCBI's gene2pubmed file, which covers every species, so it is filtered to *C. elegans* (NCBI taxon 6239) as it is downloaded.

**Curated gene sets.** The neuronal and synaptic gene sets are keyed by WormBase identifier, so they cannot drift when a gene is renamed. A laboratory substituting a gene set of its own will usually have gene names instead, so the build also accepts names and looks each up in the WormBase release it reads. A name that matches no gene is not dropped silently: it is listed in a text file written next to the database and summarized in the build log.

### Supplementary Note 4. Narrowing a search with prefixes

Typing `locomotion` into the search box returns everything that mentions the word: genes whose description does, the locomotion phenotypes and the GO terms for locomotion behavior, among the other fields shown in Supplementary Figure S1. That breadth suits a first look but not the assembly of a defined gene list. For that, WormIE offers targeted search through prefixes. A prefix confines the search to one kind of record, so `pheno:locomotion` returns only the genes with a locomotion phenotype and `expressed:AVA` only the genes CeNGEN detects in the AVA neuron type. There are seventeen prefixes: `go`, `cat` (WormCat category), `neuronal` and `syn` (the two curated gene sets), `chr` (chromosome), `domain` (protein domain), `pheno`, `tissue`, `expressed` (CeNGEN neuron type), `ortho` (ortholog), `allele`, `name`, `desc` (description), `seq` (sequence name), `wb` (WormBase identifier), `status` and `type` (gene type). For `name`, `seq`, `wb` and `desc`, a tilde in place of the colon makes the value a pattern, with `*` for any characters and `?` for one, so `name~daf-?` matches daf-2 and daf-7 alike. The `expressed` prefix also takes a threshold or a level, so `expressed:AVA@stringent` applies CeNGEN's strictest call (threshold 4 in Supplementary Note 3), `expressed:AVA>10` asks for a bulk expression above 10 and `expressed:AVA,AVB` accepts either neuron. Prefixes can also be combined with `AND`, `OR` and `NOT`, allowing layered filters. Hence `pheno:locomotion AND expressed:AVA` asks for the genes that satisfy both, a question that otherwise means searching two websites and comparing the two lists by hand. Some prefixes match a fixed field rather than text: `go`, `cat`, `neuronal`, `syn`, `status`, `type`, `wb`, `chr`, `domain` and `expressed`. When a query combines one of these with `AND`, `OR` or `NOT`, it is read left to right, so `a OR b AND c` means (a OR b) AND c. The results of such a query are not ranked by relevance.

Phenotypes are recorded under their full names in WormBase, but WormIE also recognizes the abbreviations used in the field. Searching `pheno:Vab` finds the phenotype WormBase records as *organism morphology variable* and `pheno:Him` the one recorded as *high incidence male progeny*, so a user need not know the exact wording of either.

### Supplementary Note 5. Choosing the co-expression threshold

Each gene page lists the gene's co-expression neighbors: the genes whose expression rises and falls together with it across the 52 neuron types CeNGEN covers. Co-expression is computed by correlating log-transformed expression across those 52 types. Left at that, any two genes would have a correlation value, and the 22,285 genes in CeNGEN's matrix would give about 248 million pairs. Before applying any condition, WormIE drops the genes whose expression is flat across all 52 types (a variance below $10^{-8}$ on the log scale). Every gene the variance filter removes is below 0.1 in all 52 types, the value under which Supplementary Note 3 treats a gene as not detected, so the filter trims only the genes expressed in none of the neurons. That removes 2,120 genes, about 10%, leaving 20,165 genes and about 203 million pairs. To keep only the pairs worth showing, WormIE then applies two conditions, explained in turn below: the correlation must be statistically significant, and each gene must rank among the other's 200 most correlated genes. The number of neighbors therefore varies from gene to gene.

The first condition tests each correlation with the usual $t$-test, $t = r\sqrt{n-2}/\sqrt{1-r^2}$ on $n-2$ degrees of freedom. With 203 million comparisons, the natural instinct is to correct for all of them. A pair would then need a correlation of at least 0.745 to pass, and 3,581 of the 20,165 genes would keep no neighbor at all. Worse, the genes that go empty are the narrowly expressed ones that say most about a neuron type, while broadly expressed genes keep long lists that say little. Correcting for every pair also guards against the wrong error, because a neighbor list is read from the top and a false pair far down a gene's ranking costs nothing. WormIE instead applies a p-value below 0.001 to each pair without correction, which at 52 observations means a correlation of at least 0.443 (0.273 would pass at 0.05 and 0.354 at 0.01). Only positive correlations count, so a gene whose expression runs opposite to another's is never listed as its neighbor. A p-value of 0.001 is the strictest conventional level we found that still leaves every gene a usable list.

Although the significance threshold reduces the number of partners, it does not settle the broad-versus-narrow problem above. Even at the relatively strict p-value of 0.001, the median gene has 595 neighbors while the mean is 735, indicating that a minority of genes carry very long lists. Those are the broadly expressed genes that correlate with much of the genome and so appear in nearly every neighbor list. Tightening the p-value to $10^{-6}$ reduces the median from 595 to 215 but the mean only from 735 to 380. In other words, typical genes lose most of their neighbors while the most highly connected genes retain many.

WormIE's second condition accounts for the problem by requiring the ranking to hold from both sides, a rule we call reciprocity. Each gene ranks every other gene by how strongly the two correlate, and a pair is kept only when each gene appears among the other's top 200. A broadly expressed gene correlates with thousands of genes and so earns a place in the top 200 of many of them. Those partners are rarely found in its own top 200, however, because only 200 of its thousands fit, so most of its pairs are dropped while a more specific pairing between two genes survives. Supplementary Table S2 shows how this rule, as well as the other variants considered, affects the identification of co-expression neighbors.

Reciprocity does what a stricter p-value could not. It brings the mean down from 735 to 102 and the median from 595 to 98, effectively bringing the two values together. The long lists of the broadly expressed genes are therefore mostly gone, and every gene is left with a list of comparable length. Since a gene's partners must come from its own top 200, no list can exceed 200 neighbors. The table costs 110 MB, compared with 800 MB when neighbors are selected by statistical significance alone.

In the build reported in this manuscript, the table holds 2,032,608 rows (one for each direction of a pair), or 1,016,304 pairs. Only genes that kept at least one neighbor are stored: 19,904 of the

Supplementary Table S2: **Co-expression retention under each rule.** Every rule shown also requires significance at a p-value below 0.001. Counts are exact, computed over all 20,165 genes by `tools/coexpr_rules.py` in the manuscript repository; the median and mean are over genes with at least one neighbor. Storage is the size of the co-expression table, which holds two rows per pair, scaled from the built table. The chosen rule is marked.

| **Rule** | **Median neighbors** | **Mean** | **Pairs** | **Storage** |
|---|---|---|---|---|
| significance only | 595 | 735 | 7.41M | 800 MB |
| each in the other's top 50 | 10 | 17 | 0.15M | 16 MB |
| each in the other's top 100 | 28 | 42 | 0.40M | 43 MB |
| each in the other's top 200 (used) | 98 | 102 | 1.02M | 110 MB |
| each in the other's top 400 | 243 | 237 | 2.38M | 257 MB |

20,165 in the comparison appear in the table and the other 261 had no pair that satisfied both conditions.

### Supplementary Note 6. What the network connectivity test compares against

Network connectivity asks whether the genes in a list have more recorded interactions with one another than a random list of the same size would. However, some genes are far more studied than others, so a well-studied gene carries hundreds of recorded interactions and a neglected one none. Drawing a random list of the same length from all genes would therefore make almost any list of well-studied genes look unusually connected, whether or not its members share a function. WormIE instead builds its random lists by replacing each query gene with a gene that has a similar number of recorded partners. It sorts the genes in the network by their number of partners. Walking from the fewest partners upward, it cuts that order into consecutive bands of at least 100 genes, cutting only where the count changes so that genes with the same number of partners stay together. It then swaps each query gene for a random gene from its own band (Supplementary Figure S9). A thousand such lists are drawn and the interactions within each are counted, which gives an expected count (their mean) and a spread (their standard deviation). The z-score is the distance between the query's count and that expected count, measured in units of the standard deviation.

To see how much this choice matters, we compared the two ways of building the random lists on a list of the 60 genes with the most recorded partners among the 9,099 genes and 44,191 recorded pairs in the network, a list that shares no known biology (Supplementary Table S3).

Supplementary Table S3: **The two ways of drawing random lists, tested on the best-connected genes.** The query holds the 60 genes with the highest degree in the BioGRID interaction network, where a gene's degree is its number of recorded interaction partners. Observed is the number of interactions among the 60 genes and Expected the mean over the random lists, drawn either from all genes with equal chance (uniform) or from the bands described above (degree-matched). The first pair of rows uses BioGRID 5.0.262, the release in the reported build, and the second an earlier release, as a check. Under the uniform draw the z-score moves by about ten from run to run, because the spread of the random counts is so small. For this list the degree matching is loose, since few genes have hundreds of partners: the random lists have a median of 102 partners against the query's 135.

| **Random lists** | **Observed** | **Expected** | **z** |
|---|---|---|---|
| uniform (5.0.262) | 363 | 2.0 | +144.6 |
| degree-matched (5.0.262) | 363 | 594.1 | −2.8 |
| uniform (5.0.259) | 373 | 1.9 | +155.8 |
| degree-matched (5.0.259) | 373 | 600.3 | −2.8 |

Drawn with equal chance, the random lists hold about two interactions among their members, and the 363 observed make this list look like one of the most connected sets in the genome. Matched by degree, the random lists hold about 594 interactions, more than the list itself, so the result reverses rather than merely shrinking. The reason is that in this network the genes with the most partners interact mostly with genes that have few, not with one another: across the 44,191 recorded pairs, the partner counts of the two genes in a pair are weakly negatively correlated ($r = -0.09$). A list made of the best-connected genes is therefore less connected among its members than a random list of equally well-connected genes.

For a list with real biology behind it, the matching works as intended. Take the 250 genes WormBase annotates with the phenotype aldicarb resistant: the degree-matched random lists have the same median partner count as the query, and the list still scores $z = 2.5$ against those random

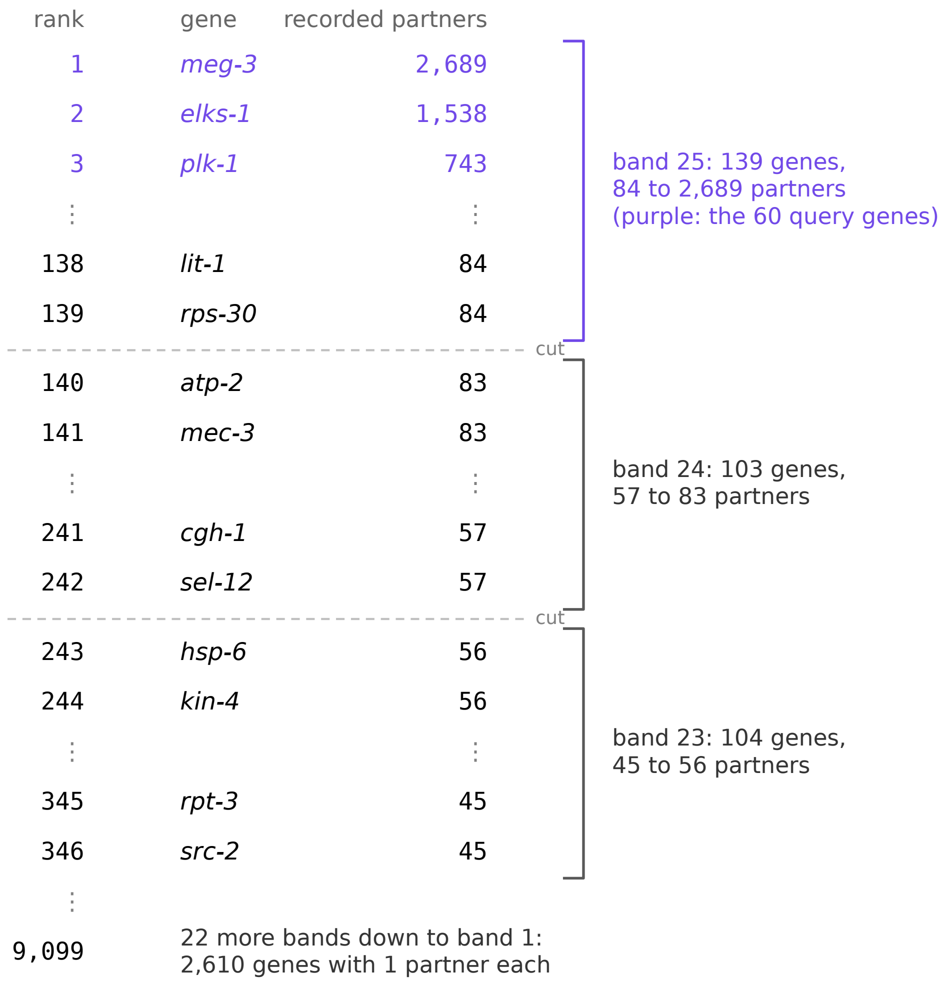


Supplementary Figure S9: **How the network connectivity test matches genes by their number of partners.** Every gene in the BioGRID network is ranked by its number of recorded partners, and the ranking is cut into consecutive bands of at least 100 genes, cutting only where the count changes so that genes with the same count stay together. A query gene is swapped for a random gene from its own band. The ranking is walked from the fewest partners upward; a band is cut where the count changes once it holds 100 genes, and the genes left over at the top join the last band closed. The list shows the first and last genes of the three highest of the 25 bands, with ties ordered by name. The top band spans 84 to 2,689 partners, and the 60 best-connected genes tested in Supplementary Table S3 (purple) all fall in it. BioGRID 5.0.262, as in the reported build, with 9,099 genes and 44,191 recorded pairs.

Alt text: A list of genes ranked by their number of recorded partners, with brackets that group the ranking into three bands, the top band in purple for the 60 query genes, and a dashed line at each cut between bands.

lists. WormIE offers both ways of drawing so that users can compare the two directly.

### Supplementary Note 7. Preparing counts and fold changes for display

Differential expression applies two transformations on the user's behalf before any result is shown, one to the counts behind the sample views and one to the fold changes. This note explains what each corrects.

#### Variance stabilization

WormIE keeps a variance-stabilized version of the counts alongside the normalized one. It exists because an abundant gene scatters more between replicates than a scarce one. The usual cure, a log transform of the normalized counts, only reverses that imbalance, since on a log scale the scarce genes scatter most (Love *et al.* 2014). An imbalance of either kind is especially detrimental for visualization. A comparison between samples pools thousands of genes at once, and whichever vary on the largest scale dominate the picture. Variance stabilization acts as a log transform on abundant genes while compressing the differences among low counts, so every gene varies on roughly the same scale.

#### Shrinking log fold changes

A gene with only a handful of reads can show a large fold change, since a few counts either way are enough to double the number. DESeq2 corrects for this by pulling each estimate toward zero in proportion to how uncertain it is, a step called shrinkage. DESeq2 offers three methods, and WormIE follows DESeq2's own rules for which one applies. apeglm, the default in both, shrinks only a comparison whose baseline is the reference condition, the group every other group is measured against. For any other comparison, including one that measures the reference against another group rather than the other way round, DESeq2 offers two methods, ashr and its original normal method, and leaves the choice to the user. WormIE selects ashr in advance for that case. The authors of apeglm find that ashr preserves large fold changes about as well as apeglm, while the normal method shrinks them too aggressively (Zhu *et al.* 2019). The user can still choose apeglm, ashr or no shrinkage on the page.

Shrinkage changes only the fold changes, so the adjusted p-values stay as DESeq2 computed them at the user's significance level. Should shrinkage fail for a comparison, the unshrunken estimates are reported instead: the page raises a warning carrying the R error and the words "showing unshrunken log2 fold changes", the contrast is labeled "shrink: none" and the methods text WormIE exports records that no shrinkage was applied.

### Supplementary Note 8. Reducing redundant enrichment terms

An enrichment table often lists several terms that say the same thing about the same genes. In GO this happens because terms nest: a list of genes that act at the synapse comes back with the synapse term, its parent and its grandparent, each on its own row with its own p-value. In the other collections it happens when two terms are carried by largely the same genes. Either way the table grows without saying more, and the distinct findings in it become harder to pick out. WormIE offers the option to reduce any enrichment table, keeping one term from each group of near-synonyms, with the procedure of REVIGO (Supek *et al.* 2011), which was designed for GO and is extended here to the other sources. This note gives the procedure and the choices made where REVIGO left them open.

#### Measuring the similarity of two GO terms

Two GO terms are scored by SimRel (Schlicker *et al.* 2006), the default of the four similarity measures REVIGO offers. SimRel asks how specific the closest ancestor shared by the two terms is. Specificity comes from how many genes carry a term: its frequency $p$ is the fraction of annotated genes carrying the term or any of its descendants, and the rarer the term, the more specific it is. The formula below measures that specificity as $-\ln p$, which is large when $p$ is small. REVIGO takes these frequencies from the UniProt GO annotations of a chosen species, whereas WormIE computes them from the same WormBase annotations it tests enrichment against, after the propagation described in Supplementary Note 2. We kept to WormBase so that the reduction runs on the same annotation set as the result it reduces, works offline and can be pinned to a WormBase release. For two terms $a$ and $b$ whose most specific shared ancestor is $c$, SimRel is

$$\mathrm{SimRel}(a,b) = \frac{2\ln p(c)}{\ln p(a) + \ln p(b)} \times (1 - p(c)).$$

The first factor is the share of the two terms' specificity that their shared ancestor already carries: 1 for identical terms, and falling as the closest shared ancestor becomes more general. The second factor pulls the score towards 0 when that ancestor is itself nearly universal, so two unrelated processes do not count as similar merely because both are biological processes. Both factors depend on which terms count as ancestors, and REVIGO does not say which relations it follows to find them. WormIE follows the same relations it propagates over (Supplementary Note 2), so that a term's frequency and its ancestors come from one hierarchy.

#### Deciding which term to keep

The procedure takes the most similar pair of terms still in the list and, if their similarity is above the cutoff, folds one of the two under the other. A folded term is not deleted: it stays in the table under the term that replaced it, with its p-value and adjusted p-value untouched, and only leaves the plots. WormIE follows REVIGO's order of rules to decide which of the two is folded:

1. a term covering more than 5% of annotated genes is folded, because a term that broad cannot stand for a cluster;

2. failing that, the term with the larger p-value is folded;

3. failing that, when one term is the parent of the other, the child is folded, unless the child makes up more than 75% of the parent, in which case the parent is folded;

4. any remaining tie is broken by keeping the term with the smaller GO identifier, the older of the two, instead of at random as in REVIGO, to ensure that the same list always reduces the same way.

This repeats until no remaining pair is more similar than the cutoff. The cutoff defaults to REVIGO's own 0.7 and can be moved anywhere between 0 and 1 by the user, with a lower cutoff folding more terms. Each of the three GO aspects is reduced on its own. In ranked-list enrichment a set with a positive enrichment score is never compared with one with a negative score, since the two describe opposite changes even when their genes overlap. Scoring every pair of terms against every other grows quickly with the length of the list, so a list of more than 2,000 terms is left unreduced until the adjusted p-value cutoff is tightened.

#### Annotations without a hierarchy

The other collections carry no hierarchy that WormIE can score, so the same procedure runs on the genes behind each term instead. Similarity is scored by gene overlap: the share of genes the two terms have in common out of all the genes in either (the Jaccard index). In over-representation these are the query genes each term holds; in ranked-list enrichment they are each set's leading edge, the genes that carried its score. A term is treated as general when it covers more than 5% of the genes tested, and when one term's genes include all of another's, the larger term is treated as the parent. We calibrated the gene overlap cutoff against SimRel on the GO enrichment of the four gene lists of Supplementary Note 11 (WormIE and published, up and down), choosing the value that shortens the lists by about as much as SimRel does at 0.7, which gave a default of 0.5 (Supplementary Table S4).

Supplementary Table S4: **Terms kept by the two reductions on the GO lists of the case study.** Each list holds the GO terms enriched at an adjusted p-value of 0.05 or below across the three GO aspects, from the lists of Supplementary Note 11. The remaining columns give the terms kept after reduction by SimRel at its default cutoff of 0.7 and by gene overlap at three cutoffs. The shaded columns are the two defaults.

| | | **SimRel** | **Gene overlap** | | |
|---|---|---|---|---|---|
| **Gene list** | **Enriched terms** | **0.7** | **0.3** | **0.5** | **0.7** |
| WormIE, up-regulated | 299 | 93 | 64 | 110 | 145 |
| WormIE, down-regulated | 197 | 107 | 54 | 86 | 125 |
| Published, up-regulated | 268 | 82 | 55 | 98 | 124 |
| Published, down-regulated | 195 | 107 | 53 | 84 | 123 |

### Supplementary Note 9. Reading per-sample gene set scores

Per-sample gene set scoring answers a question that a two-group comparison cannot: in a series such as adults profiled at day 1, 3, 5, 7 and 10, which gene sets rise or fall steadily with age? Differential expression would give ten pairwise gene lists and no trend, whereas scoring each sample on its own gives every gene set one number per sample, a trajectory with the replicates still visible. The scores follow the ssGSEA method (Barbie *et al.* 2009) as implemented by GSVA (Hänzelmann *et al.* 2013), including the scaling that GSVA applies by default. For each sample, the genes are ranked from the most to the least expressed, and a gene set's score measures how far its members crowd toward the top or bottom of that ranking. Within that score, genes near the top of the ranking are weighted a little more than the rest, by GSVA's default exponent of 0.25, making a gene's position matter without letting the handful of most expressed genes dominate. The score is near zero when the set's genes are scattered evenly through the ranking, high when they sit near the top and low when they sit near the bottom. Once every score is computed, GSVA divides each one by the range between the largest and the smallest score in the table. This scaling makes scores comparable within a run, but it also limits comparisons between runs. The divisor belongs to the run rather than to any one sample, so adding a sample or a gene set changes the divisor and every score in the table. Scores should therefore be compared only among samples scored in the same run, never between separate runs, even when the same gene sets were used.

One default in WormIE differs from GSVA's. A built-in set is scored only when 15 to 500 of its genes are present in the matrix, the default bounds of the original GSEA software, where GSVA sets none. Meanwhile, a collection the user supplies as a GMT file is scored without a bound.

The output is a table with one row per gene set and one column per sample. Any two scores in it can be compared, but no score comes with a statistical test attached. To claim a trend across a series, WormIE fits each gene set's scores across the samples against a variable that describes the samples. The variable can be continuous, such as age or dose, or categorical (a factor, in R's terms), such as genotype. A test needs an estimate of noise, which the two kinds obtain differently.

For a continuous variable, WormIE fits a straight line by default, which tests for a steady rise or fall. When the variable takes five or more distinct values, such as five timepoints, WormIE can fit a curve instead. Either way, the fit supplies the noise estimate itself, because the noise is how far the points fall from the fitted line or curve. Five single samples at five ages are therefore enough, and three samples are the minimum.

For a categorical variable, the test asks whether the levels differ at all, for example whether mutant and wild type have different scores. Unlike a continuous variable, a categorical one gives no line to fit: each level's estimate is simply the average of its own samples, so replicates are the only source of a noise estimate. With one sample per level there is nothing to compare within a level, and WormIE does not run the fit but suggests treating the variable as continuous where it is a number, such as timepoints.

Whichever the variable, a few samples give a poor estimate of how much a set's scores vary. The fit therefore pulls each set's estimate toward the typical value across all sets, the moderation limma introduced for genes (Ritchie *et al.* 2015), which steadies the test for every set. This is how the authors of GSVA recommend testing per-sample scores (Hänzelmann *et al.* 2013). The test runs in the browser in a second R session, started from the same R bundle as differential expression with limma added, so it pays its own start-up wait. The result table gives each set's slope, its difference between two levels or its F statistic across several, with its p-value and its Benjamini-Hochberg adjusted p-value.

### Supplementary Note 10. Timings and agreement with the original packages in native R

The main text reports that the analyses WormIE reimplements agree with the R packages they follow. This note gives the measurements behind that claim, the machine they were taken on and how long each analysis takes.

#### What was measured

Each analysis was run on the same input in WormIE and in R, and the two outputs were compared set by set. One setting was changed from WormIE's default: over-representation normally skips a set that shares fewer than two genes with the query list, and the benchmark lowered that to one so that every set with a gene in common was tested in both tools. Because the inputs are synthetic and generated from a fixed seed, anyone can recreate the same inputs. Agreement on real data is shown separately by Supplementary Note 11, which repeats a published analysis from reads to enriched categories. Supplementary Table S5 lists the four analyses benchmarked, with the input each one takes and the signal planted in each input. The planted signal makes some sets come out significant, so the two pipelines have calls to agree on. Per-sample gene set scoring carries no planted signal, because they are compared on the scores themselves. Where the main text quotes a count of significant sets, it is the total across the configurations of Supplementary Table S6. The ranked-list counts in that table come before refinement, and refinement brings them to the 114 of 114 the main text reports, as described below.

Supplementary Table S5: **What each benchmarked analysis was given.** The three gene set analyses share the same collections of gene sets, drawn from the same gene pool as the ranked list.

| Analysis | Input | Planted signal |
|---|---|---|
| Differential expression | count table | 10% of the genes carry a specified fold change |
| Ranked-list enrichment | ranked gene list, gene sets | 10% of the gene sets are built from genes near the top of the list |
| Over-representation | query gene list, gene sets | three of those sets are over-represented in the query list |
| Per-sample gene set scoring | expression matrix, gene sets | none |

In this benchmark, the gene sets each list was tested against were generated beforehand and handed to both tools, so any disagreement can be attributed to the test itself. Users, however, will usually test their gene lists against the built-in sets, such as Gene Ontology terms. WormIE assigns genes to GO terms by the propagation described in Supplementary Note 2 while clusterProfiler reads the Bioconductor annotation package, so the two can call different terms on the same gene list even with identical arithmetic. Whether they do is checked in Supplementary Note 2, where each tool tests one real gene list for enriched GO terms. Most results agree, and the disagreements trace mostly to the two reading different annotation releases and otherwise to terms that sit at the significance threshold.

Measurements were taken on an Apple M2 with eight cores and 24 GB of memory running macOS 26.6, with Chrome 148 for the browser arm. The R arm ran R 4.6.1 with DESeq2 1.52.0 and apeglm 1.34.0, the same package releases WormIE carries in the browser on R 4.5.1, so the comparison isolates the difference between running in R and running in the browser, and it used clusterProfiler 4.20.3, fgsea 1.38.0 and GSVA 2.6.10 for the gene set analyses. The browser runs

R 4.5.1 rather than the current release because webR, the build of R for browsers that WormIE uses, provides R 4.5.1. Every number in Supplementary Tables S6 and S7 comes from runs in October 2026 on an otherwise idle machine, with R's random number generator seeded so that the permutation-based results repeat exactly. Values were compared at full precision rather than at the digits printed, so rounding hides no difference. No log fold change differs by more than $1.8 \times 10^{-7}$ between the two runs and no adjusted p-value by more than $4 \times 10^{-14}$, the residue of compiling the package to run in the browser and nothing else. Timings are the mean over ten runs, except the WormIE differential expression column, which is the mean of three.

Supplementary Table S6: **Timing and agreement for analyses of synthetically generated data.** The correlation is on log fold changes for differential expression, on normalized enrichment scores for ranked-list enrichment, on $-\log_{10}$ of the p-value for over-representation and on the scores themselves for per-sample gene set scoring. Sign flips are counted among genes with an adjusted p-value below 0.05. Set counts are the sets R calls significant at an adjusted p-value below 0.05 that WormIE also calls, followed by those WormIE adds. Universe is the number of genes the sets are drawn from.

| **Analysis** | **Input** | **R** | **WormIE** | **Agreement** |
|---|---|---|---|---|
| Differential expression | 1,500 genes, 6 samples | 0.7 s | 4.4 s | $r = 1.0000$, no sign flips |
| | 8,000 genes, 6 samples | 2.0 s | 10.6 s | $r = 1.0000$, no sign flips |
| | 18,000 genes, 8 samples | 4.2 s | 20.5 s | $r = 1.0000$, no sign flips |
| Ranked-list enrichment | 2,000 universe, 50 sets | 135 ms | 25 ms | $r = 1.0000$, 5 of 5, none added |
| | 2,000 universe, 500 sets | 173 ms | 40 ms | $r = 1.0000$, 51 of 52, 1 added |
| | 10,000 universe, 50 sets | 122 ms | 25 ms | $r = 1.0000$, 5 of 5, none added |
| | 10,000 universe, 500 sets | 186 ms | 42 ms | $r = 0.9999$, 51 of 51, none added |
| Over-representation | 2,000 universe, 50 sets, 100 genes | 8 ms | 2 ms | $r = 1.0000$, 3 of 3, none added |
| | 2,000 universe, 50 sets, 500 genes | 8 ms | 2 ms | $r = 1.0000$, 4 of 4, none added |
| | 2,000 universe, 500 sets, 100 genes | 46 ms | 16 ms | $r = 1.0000$, 37 of 37, none added |
| | 2,000 universe, 500 sets, 500 genes | 51 ms | 16 ms | $r = 1.0000$, 41 of 41, none added |
| | 10,000 universe, 50 sets, 100 genes | 11 ms | 1 ms | $r = 1.0000$, 3 of 3, none added |
| | 10,000 universe, 50 sets, 500 genes | 11 ms | 2 ms | $r = 1.0000$, 3 of 3, none added |
| | 10,000 universe, 500 sets, 100 genes | 142 ms | 13 ms | $r = 1.0000$, 6 of 6, none added |
| | 10,000 universe, 500 sets, 500 genes | 146 ms | 20 ms | $r = 1.0000$, 24 of 24, none added |
| Per-sample gene set scoring | 2,000 universe, 50 sets | 29 ms | 14 ms | $r = 1.0000$, max $\Delta$ 0.0001 |
| | 2,000 universe, 500 sets | 69 ms | 51 ms | $r = 1.0000$, max $\Delta$ 0.0001 |
| | 10,000 universe, 50 sets | 39 ms | 76 ms | $r = 1.0000$, max $\Delta$ 0.0000 |
| | 10,000 universe, 500 sets | 87 ms | 216 ms | $r = 1.0000$, max $\Delta$ 0.0001 |

Supplementary Table S6 gives the timing and agreement for every input size that was run. Differential expression is the one analysis that runs R itself inside the browser, and it takes about five times as long there as in native R. The three gene set analyses, rewritten for the browser, finish within a few hundred milliseconds, at worst about two and a half times native R's time and often faster, because the gene sets are already indexed in the local database.

Comparing the two runs side by side yields two disagreements, both in ranked-list enrichment and both tracing back to a set sitting at the significance threshold. The over-representation runs agree on every set: the 1,918 p-values match to within $4 \times 10^{-11}$ and the adjusted p-values to within the same, since both tools drop the sets that share no gene with the query list before the Benjamini-Hochberg step.

Both sets come from the input with a universe of 2,000 genes and 500 sets. The two tools agree on the sets' enrichment scores (1.86 and 1.65) yet disagree on their p-values. That is expected, because the p-value is the share of 1,000 random gene sets that score at least as high as the real one, so a few random sets more or fewer can move a set across the threshold. The first set is lost in WormIE and significant in fgsea only, because its raw p-value of 0.007 in WormIE and 0.002 in fgsea adjusts to 0.063 and 0.022. The second is an addition found in WormIE only, because its raw p-value of 0.005 in WormIE and 0.011 in fgsea adjusts to 0.044 and 0.097. Part of this gap may come from WormIE's decision to let sets of similar size share one collection of random scores, as described in the main text. Even so, the cost is small here, since the scores still correlate at 0.9999 or better and these two sets are the only ones whose calls differ.

Besides the trouble with sets that hug the threshold, reshuffling brings a second problem. With 1,000 reshuffles no p-value can fall below 1 in 1,000, so the strongest sets all stop at that limit and cannot be ranked against each other. Across the four inputs, 80 sets had a p-value within three times that limit, close enough that one or two random sets decide the p-value. For sets that strong, WormIE refines the estimate the way fgsea does, by continuing to reshuffle among ever more extreme outcomes until the p-value is pinned down, to a floor of $10^{-50}$. The 80 refined sets agree with fgsea's refined estimates at $r > 0.998$ on $-\log_{10} p$. Both tools place 32 sets at the $10^{-50}$ floor, and one further set reaches it in WormIE while fgsea leaves it just above, at $6.5 \times 10^{-50}$. Refinement also touches the two sets the tools disagreed on and settles both: the first moves to an adjusted p-value of 0.008 in WormIE and 0.010 in fgsea, the second to 0.037 and 0.029. WormIE therefore recovers all 114 sets fgsea calls significant and adds none. Because a single refined estimate is unstable at this depth, we ran fgsea eight times on the 36 refined sets of the input with a universe of 2,000 genes and 500 sets, changing only the random seed. Between runs a set's p-value varied by a factor of 13 at the median and by more than 100 at worst, for a set near $10^{-46}$. Averaged over eight runs of each tool on those sets, WormIE's p-values sit at 1.1 times fgsea's, well within that spread.

#### The cost of starting R in the browser

The gene set analyses were reimplemented rather than run from their original packages. Running the originals would have meant fetching each package and starting R inside the browser first, a wait of seconds before any analysis could begin, or longer on a slow connection. The three tests are arithmetic over gene sets, counting how many of a list's genes carry a term or walking down a ranked list, and those sets already sit on the machine, so implementing the tests directly avoids that wait. The original packages were never loaded in the browser, so the wait is known only for DESeq2, the one package WormIE does load (Supplementary Table S7).

Supplementary Table S7: **What it costs to start R in the browser.** Each value is the mean ± standard deviation of ten runs. Bundle is the download of the R packages, read from the same machine here, so it is a lower bound on what a visitor pays over a network; starting R and loading DESeq2 are computation and carry over to any machine.

| **Setting** | **Total** | **Bundle** | **Start R** | **Load DESeq2** |
|---|---:|---:|---:|---:|
| First visit, nothing cached | $15.5 \pm 0.1$ s | $0.3 \pm 0.0$ s | $0.6 \pm 0.0$ s | $14.2 \pm 0.1$ s |
| Return visit, bundle cached | $15.0 \pm 0.1$ s | $0.1 \pm 0.0$ s | $0.6 \pm 0.0$ s | $14.1 \pm 0.1$ s |
| One further instance in the same page | $10.4 \pm 0.1$ s | 0.0 s | $0.3 \pm 0.0$ s | $9.7 \pm 0.0$ s |

Loading DESeq2 dominates every setting, and caching the packages does not shorten that step. A user who opens the differential expression page therefore waits about 15 s before any model is fitted, and each further copy of R started to extract contrasts in parallel adds about 10 s more. The 100 MB of packages also has to be fetched once over the network, which the table does not include. Loading DESeq2 runs on a single thread and is sensitive to other load on the computer: on the same machine it took about 14 s when idle and around 40 s with other work competing for the processor, so a user on a busy laptop should expect a longer wait than the table shows. The wait for the three gene set packages would be of the same order or longer, since loading time grows with the number of packages pulled in and fgsea, GSVA and clusterProfiler depend on 31, 88 and 120 other packages where DESeq2 depends on 45. Ranked-list enrichment itself runs in 25 to 42 ms (Supplementary Table S6), so reimplementing the three analyses avoids a startup cost two to three orders of magnitude larger than the analyses themselves.

### Supplementary Note 11. Reproducing a published analysis

The main text's section *A published analysis, reproduced in a browser* repeats the germline RNA-seq analysis of Robert *et al.* (2023) in WormIE. This note lists the published pipeline and WormIE's side by side, step by step, and reports how closely the two agree at each step, with the tables the main text summarizes.

The Robert *et al.* (2023) analysis is a good test for two reasons. First, it follows the route of the main text's *In-browser analysis* section from end to end: counting reads, fitting a differential expression model and testing the result for enriched categories. Second, every input is public, so the comparison rests on the authors' data rather than on the paper's description of the data.

We ran the analysis on database build 20261025-112346 and repeated it on the later build 20261026-105834, which gave identical tables at every step.

#### The two pipelines

Supplementary Table S8 lists every step of both pipelines, and the text below explains each step where the two differ.

The pipeline starts from raw sequencing reads, which have to be trimmed and aligned before anything can be counted. WormIE does not trim or align reads, so we ran both steps outside WormIE with the authors' tools and versions, fastp 0.20.1 and STAR 2.7.3a. Counting the aligned reads is the first step both pipelines share, and it needs gene models to count against. WormIE's build carries the gene models of WormBase WS298 and uses them by default, but the authors aligned to the earlier release WS278, so we supplied the WS278 gene models as a GTF file in place of WormIE's own. The paper does not specify the strandedness for counting, so we went with WormIE's default, which is unstranded. A check after counting showed that our totals match the authors' deposited counts to within 0.15% on every library. The counting program itself could not be matched, since the authors used htseq-count and WormIE uses featureCounts. To count inside the browser, WormIE needs a program that compiles to WebAssembly directly, and featureCounts does because it is written in C, while htseq-count is a Python program and would need a Python interpreter in the browser as well.

The step after counting is differential expression, and both pipelines fit the same model: genotype plus the day each replicate was prepared. The day term is not stated in the authors' Methods but comes from the R script those Methods cite. Beyond the model, three settings in that script had to be matched. It leaves the significance level, which decides how many low-count genes DESeq2's independent filtering sets aside before the p-values are adjusted, at DESeq2's default of 0.1, and WormIE kept that default. It does not shrink fold changes, whereas WormIE does by default with apeglm, so shrinkage was switched off. It drops low-count genes only after the fit, in the post-fit low-count filter, so WormIE's pre-filter, which drops genes with fewer than 10 reads in total before the fit, was switched off to match.

The authors, and therefore this comparison, then filtered the results before building the gene lists to keep only nuclear protein-coding genes with a mean normalized count (DESeq2's baseMean) above 10. We call this the expression filter. WormIE follows the DESeq2 convention and has no filtering step after the fit, so the expression filter was applied outside WormIE. The expression filter also removes every gene the pre-filter would have removed, because a gene with fewer than 10 reads in total never reaches a mean normalized count above 10. Switching the pre-filter off therefore removes no gene from the comparison. We also ran the fit once with the pre-filter left on, as WormIE does by default, and report that run only here. It changes the lists only slightly, since WormIE then calls 399 genes up and 679 down and recovers 381 of the 387 published up and 661 of the 663 published down genes, against 383 and 661 with the pre-filter off. The expression filter itself also matters little for this dataset. Without the expression filter WormIE calls 428 genes up and 685 down, against 394 and 664 with the filter. The difference is about five percent, so a user who skips the filter should get nearly the same lists.

Supplementary Table S8: **The published pipeline versus the WormIE run for the analyses of the Robert *et al.* (2023) transcriptomic data.**

| **Step** | **Published** | **WormIE** |
|---|---|---|
| Trimming | fastp 0.20.1 | fastp 0.20.1 [outside WormIE ] |
| Alignment | RNA-STAR 2.7.3a, WS278 | RNA-STAR 2.7.3a, WS278 [outside WormIE ] |
| Counting | htseq-count 0.7.2, WS278, unstranded | featureCounts, subread 2.0.6, WS278, unstranded |
| Expression filter | nuclear protein-coding, mean normalized count > 10 | the same [outside WormIE ] |
| Differential expression | DESeq2 1.36.0, R 4.2.2 | DESeq2 1.52.0, R 4.5.1 [in the browser] |
| Model | genotype + day of preparation | the same [day of preparation in the Batch covariate column] |
| Post-fit low-count filter | rows under 10 reads, after the fit | none |
| Shrinkage | none | none |
| Category enrichment | WormCat | the same [Over-representation > WormCat, lists pasted] |
| Germline classification | hypergeometric test in R | the same [Custom gene set testing, one set per class] |

Throughout this note, WormIE's results are compared with the full results table the authors deposited with the paper. Filtering that table as their gene-list script does leaves exactly the 7,555 genes the paper reports, with 387 up and 663 down. Those are the numbers behind the recovery figures in the main text and the totals beside the bars in Figure 3a of the main text. The same filter on WormIE's table leaves 7,569 genes, 7,553 of them shared with the authors', so the two runs test nearly the same list and any disagreement below is between the pipelines rather than the gene lists.

#### Agreement at each step

Supplementary Table S9: **Gene-level agreement.** Gene sets are compared after the expression filter, and the last column is the Jaccard index. The WormCat rows compare the categories enriched in WormIE's own lists with those enriched when the authors' lists are run through WormIE.

| **Comparison** | **WormIE** | **published** | **shared** | **Jaccard** |
|---|---|---|---|---|
| Filtered genes | 7,569 | 7,555 | 7,553 | 0.998 |
| Called up | 394 | 387 | 383 | 0.962 |
| Called down | 664 | 663 | 661 | 0.992 |
| WormCat terms, up | 53 | 54 | 51 | 0.911 |
| WormCat terms, down | 88 | 88 | 88 | 1.000 |

Per-library counts agree at a Pearson correlation of 0.9988 or better on every one of the six libraries (three wild type and three *sin-3(syb2172)*), computed on $\log_{10}$ of the count plus one so that a few highly expressed genes cannot carry the correlation. Library totals differ by between 0.10 and 0.14 percent. No gene is called significant in opposite directions by the two runs. At an adjusted p-value below 0.05, the full WormIE run, from its own counts, calls 394 genes up and

664 down against the published 387 and 663 (Supplementary Table S9). To separate the counting step from the fit, we also ran WormIE's DESeq2 on the authors' deposited counts, which recovers 386 of their 387 up and 662 of their 663 down genes and calls no others. The authors' model run in native R with DESeq2 1.52.0 on the same counts reproduces their calls exactly, so the two DESeq2 releases agree. The one gene lost in each direction most likely reflects the authors' post-fit low-count filter, and the rest of the difference in the full run would then come from counting with featureCounts rather than htseq-count. Summed over both directions the WormIE run calls 1,058 genes significant and the published one 1,050, sharing 1,044 of them, so 1,064 genes are called by one run or the other. Fitted on its own counts without the day of preparation but otherwise identically, WormIE recovers only 92% and 89% of the published calls, so leaving that term out costs far more agreement than any difference between the pipelines.

#### WormCat enrichment

In WormIE, the natural next step after differential expression is to hand the results over to over-representation, which takes the up and down lists, or to ranked-list enrichment, which takes every gene ranked by its test statistic. A list handed over to over-representation this way is tested against every gene in the contrast's results table. WormIE chooses this background by default, on the reasoning that only the genes tested in the contrast could have been called differentially expressed. For their Figure 3C, Robert *et al.* (2023) tested the up and down lists by over-representation against WormCat's whole-genome annotation instead, as WormCat's own tool does. To match them, we pasted the two lists as plain gene lists, which WormIE tests against all 31,389 genes of that annotation. Running the authors' own gene lists through WormIE gives lists of enriched categories that overlap those from WormIE's own lists in 51 of 54 categories up and in every category down (Supplementary Table S9). The five categories up that only one run calls all sit near the cutoff, at adjusted p-values from 0.021 to 0.049, and each rests on two to seven genes. What disagreement remains between individual genes has therefore largely disappeared by the time it reaches categories.

#### Ranked-list enrichment

The published analysis tested only the up and down gene lists, so we also took the other route WormIE offers after differential expression. Every gene that passed the expression filter was ranked by its DESeq2 Wald statistic, 7,569 genes from WormIE's fit and 7,555 from the authors' table, and each ranked list was tested against WormCat's categories by ranked-list enrichment. The set-size limit was raised from the default of 15 to 500 genes to 15 to 2,000, because two of the categories the paper reports, Metabolism and Signaling, hold 1,020 and 586 of the ranked genes and would otherwise not be tested.

The two fits agree closely, and Figure 3d of the main text plots the score each gives every category. Of the 168 categories tested, 30 are significant at an adjusted p-value of 0.05 or below in WormIE's fit and 31 in the authors', and the 28 they share all move in the same direction. The five categories that only one fit calls have adjusted p-values from 0.025 to 0.084 in both fits, so they sit at the cutoff rather than mark a difference between the fits. Ranked-list enrichment calls four of the seven categories the paper reports among the down-regulated genes in both fits: ribosome, metabolism, mRNA functions and chromatin-associated transcription. It calls cell cycle, chaperone and proteolysis proteasome in neither fit, although over-representation calls all three in both. In those three a subset of genes changes sharply while the category as a whole does not shift, which over-representation detects and a ranked test does not. Of the two categories the paper reports among the up-regulated genes, the ranked test calls signaling in both fits, while it scores metabolism, which the paper reports in both directions, only among the down-regulated genes.

The ranked test also calls three categories in both fits, in the same direction, that over-representation called in neither: *Extracellular material: collagen* (up, 40 genes), *Non-coding RNA: tRNA* (down, 82 genes) and *Transcription factor: NHR* (up, 54 genes). In each, many genes change a little, each below the cutoff that defines the gene lists, so these are what the ranked method adds rather than results the published analysis missed. All three are among the categories that Ranked-list consensus keeps from the two fits, with nearly the same leading-edge genes in both (Jaccard index 0.91 to 1.00), and all three stay significant at the default ceiling of 500.

#### The germline classification

Figure 3D of the published analysis uses a hypergeometric test in R to ask whether the misregulated genes are drawn disproportionately from the oogenic or the spermatogenic classes of Ortiz *et al.* (2014). WormIE has no built-in copy of that classification, so it was supplied as custom gene sets. Restricted to protein-coding genes, it holds 1,653 oogenic, 2,412 spermatogenic and 5,923 sex-neutral genes, against the 1,660, 2,453 and 5,950 the published analysis reports, and all 10,754 of its identifiers still resolve at WS298. With no background supplied, WormIE takes every gene in the supplied sets as the background, as clusterProfiler does. Here that is the 9,988 classified genes, each in exactly one class, which is the exact background the authors used.

Supplementary Table S10: **Germline classification of the differentially expressed genes.** Genes outside the classification are dropped before testing, which is why the tested list is smaller than the called list. Adjusted p-values are Benjamini-Hochberg across the three sets.

| **List** | **Set** | **genes** | **tested** | **fold** | **adjusted $p$** |
|---|---|---|---|---|---|
| WormIE, up | oogenic | 110 | 365 | 1.82 | $1.1 \times 10^{-10}$ |
| | spermatogenic | 77 | 365 | 0.87 | 1.00 |
| | sex-neutral | 178 | 365 | 0.82 | 1.00 |
| published list, up | oogenic | 107 | 359 | 1.80 | $4.2 \times 10^{-10}$ |
| | spermatogenic | 79 | 359 | 0.91 | 1.00 |
| | sex-neutral | 173 | 359 | 0.81 | 1.00 |
| WormIE, down | sex-neutral | 479 | 614 | 1.32 | $1.2 \times 10^{-23}$ |
| | oogenic | 99 | 614 | 0.97 | 0.95 |
| | spermatogenic | 36 | 614 | 0.24 | 1.00 |
| published list, down | sex-neutral | 478 | 613 | 1.31 | $1.6 \times 10^{-23}$ |
| | oogenic | 99 | 613 | 0.98 | 0.94 |
| | spermatogenic | 36 | 613 | 0.24 | 1.00 |

The published analysis counts 108 oogenic genes among those upregulated (Supplementary Table S10). Rerunning its gene list here counts 107, one gene fewer in a classification reconstructed a decade after it was published, and WormIE's own list counts 110. Both runs put the signal in the same set, and neither finds the spermatogenic or the sex-neutral set enriched among the genes up in the mutant.
